# Developmental plasticity and genetic selection shaped cereal evolution in the Early Holocene southern Levant

**DOI:** 10.1101/2024.08.18.608467

**Authors:** Jade Whitlam, Pascal Flohr, Amy Bogaard, Mike Charles, Bill Finlayson, Cheryl A. Makarewicz

## Abstract

The domestication of plants in southwest Asia was an evolutionary process that took place over several millennia in the Early Holocene. During this time domestic species developed distinct traits that distinguish them from their wild counterparts. Current models of plant domestication emphasise the role of genetic selection in the evolution of these traits, viewing these as heritable adaptations that arose in response to selective pressures associated with human cultivation. In cereals, domestication resulted in the evolution of non-shattering rachis and increased grain size, two traits that can be tracked directly in the archaeobotanical record. Measurements of cereal grains from Early Neolithic sites indicate that grain size increase occurred prior to the evolution of non-shattering rachis, and it has been proposed that this reflects selection for larger grains under tillage, signifying pre-domestication cultivation. Here we combine morphological and metrical analysis of cereal remains, stable carbon isotope analysis, and weed ecology to test this hypothesis, using three assemblages from the southern Levant: Pre-Pottery Neolithic A Sharara (c. 9250-9200 cal BCE), Pre-Pottery Neolithic A el-Hemmeh (c. 9400-8700 cal BCE) and Late Pre-Pottery Neolithic B el-Hemmeh (c. 7500-7000 cal BCE). Our findings indicate that increased grain size in the Early Holocene is better understood as a plastic response to variation in growing conditions (specifically moisture), rather than a result of genetic selection for increased grain size under cultivation (i.e., tillage). We argue that cereal evolution in southwest Asia was initially driven by developmental plasticity, followed by genetic selection.

**Significance Statement:** Cereals were amongst the earliest crops domesticated in southwest Asia and remain central to modern agriculture. We can track cereal domestication archaeologically using a suite of morphological traits, which distinguish domestic species from their wild counterparts. The process by which these traits evolved is still not fully understood but has widely been framed as genetic adaptation to cultivation. Here we seek to disentangle how two key domestication traits relating to seed dispersal and size evolved. By taking a multi-stranded approach to new archaeobotanical evidence, we demonstrate that in the Early Holocene southern Levant, cereal domestication occurred in a sequence where developmental plasticity preceded genetic selection.

## Introduction

Plant domestication is a co-evolutionary process that occurs when wild species adapt to human-constructed environments and the novel selection pressures they impose. While the adaptive traits selected for during domestication can vary across species, convergent evolution has led to multiple crop species acquiring similar traits, known collectively as the ‘domestication syndrome’ (Hammer, 1984). For cereal crops, domestication syndrome traits include loss of shattering (indehiscence), increased grain size, loss of dispersal aids (e.g., awns, hairs), reduced seed dormancy and apical dominance (Fuller, 2007; Hammer, 1984; Harlan et al., 1973; Zohary, 1989). Loss of shattering and increased grain size have been used as key traits for domestication studies, as they can be tracked directly in the archaeobotanical record.

A necessary precondition for the domestication of crops was the cultivation of their wild progenitors, termed ‘pre-domestication cultivation’ (PDC). PDC has been inferred at a number of Early Holocene sites across the Fertile Crescent based on multiple lines of evidence, including the presence of domestic-sized grains and arable weeds that are considered indicative of a cultivated environment (Colledge, 1998; Colledge et al., 2018; Edwards et al., 2004; Hillman et al., 2001; Kislev, 1997; Riehl et al., 2013; van Zeist and de Roller, 1994; Weiss et al., 2006; White and Makarewicz, 2012; Willcox et al., 2008). It was PDC and its associated practices that created the selective environments in which domestic traits evolved. For example, in domestic barley (*Hordeum vulgare*), loss of shattering is known to have involved mutations in two adjacent complementary genes (Pourkheirandish et al., 2015). Selection for these non-shattering mutants is linked to systematic sickle harvesting and annual re-sowing, two practices that would have conferred an evolutionary advantage on plants that retained their seeds when ripe (Fuller et al., 2014, 2010; Harlan et al., 1973; Hillman and Davies, 1990; Maeda et al., 2016). In contrast, the genetic mechanisms underlying the evolution of increased grain size, and the practices that selected for this trait are not well-understood. Despite this, increased grain size features prominently in models of PDC (Colledge et al., 2018; Weiss et al., 2006; White and Makarewicz, 2012; Willcox, 2004) and the presence of ‘domestic’ or ‘cultivated-type’ cereal grains in combination with shattering (wild) rachis morphotypes at sites has been argued to represent a stage of “semi-domestication” (Fuller, 2007). Increased grain size in this context is often linked explicitly to tillage, or to general improvements in growing conditions brought about by cultivation (Colledge et al., 2018; Fuller, 2007; Harlan et al., 1973). Tillage is hypothesised to have played a role in selection for increased grain size as it creates deeper burial conditions in which larger seeds would have a competitive advantage (Harlan et al., 1973; Westoby et al., 1996). However, experimental studies have failed to support this theory (Kluyver et al., 2013). Comparable scales of seed enlargement are also observed in domesticated vegetable crops which are not propagated from seed, suggesting that indirect selection may have driven the evolution of this trait (Kluyver et al., 2017).

Studies of the evolution of increasing grain size are further complicated by the fact that grain size is a highly plastic trait. Developmental plasticity refers to the ability of an individual genotype to alter its developmental processes and phenotypic outcomes in response to different environmental conditions (Moczek et al., 2011; Zeder, 2018). This allows phenotypic adaptation to occur in the absence of genetic change. In recent years, interest in developmental plasticity as a novel evolutionary process has increased, with proponents of the extended evolutionary synthesis (EES) regarding plasticity as one of several processes that shape evolution by generating and directing variation on which selection operates (Laland et al., 2014; Müller, 2017; Pigliucci, 2007; West-Eberhard, 2003; Zeder, 2018, 2017). The EES also contends that persistent plastic traits can be transformed to become fixed components of an organism’s genome through a process of genetic accommodation (Moczek et al., 2011; Ng and Kinjo, 2023; Pfennig et al., 2010; West-Eberhard, 2003). In other words, phenotypic change may be a *cause* rather than a consequence of genetic change. While origins of agriculture and domestication studies have been quick to embrace some aspects of the EES (e.g., Niche Construction Theory) (Fuller and Stevens, 2017; Ramsey, 2023; Rowley-Conwy and Layton, 2011; Smith, 2016, 2011, 2007; Zeder, 2016, 2015), less attention has been paid to exploring plasticity and its role in plant domestication.

Here we report on evidence from two early Holocene sites in the southern Levant, which supports the hypothesis that cereal evolution was shaped by developmental plasticity as well as genetic selection, and that increased grain size is not a direct function of PDC as traditionally modelled.

In southwest Asia, where wheat (*Triticum* spp.) and barley were among the first plants to be domesticated in the Early Holocene (*c*. 9700-6000 cal BCE), archaeobotanical and genomic data demonstrate that domestication was a multicentric and protracted process (Arranz-Otaegui et al., 2016; Fuller, 2007; Fuller et al., 2012b; Pourkheirandish et al., 2015; Tanno and Willcox, 2006), involving rates of evolution comparable to those observed under natural selection (Fuller et al., 2012a; Purugganan and Fuller, 2011). Archaeobotanical data also indicate that the full suite of domestication syndrome traits did not emerge simultaneously in cereals but were subject to changing rates of evolution over time and between different regions (Allaby et al., 2017; Fuller, 2007). Increased grain size appears to have preceded the development of indehiscence, with ‘domestic’ or ‘cultivated-type’ grains reported from numerous PPNA (9700-8700 cal BCE) sites (Arranz-Otaegui et al., 2016; Fuller, 2007). Selection for loss of shattering is evidenced in the Northern Levant during the Early Pre-Pottery Neolithic B (PPNB, 8700-8200 cal BCE), based on increased proportions of non-shattering rachis morphotypes in archaeobotanical assemblages (Arranz-Otaegui et al., 2016; Tanno and Willcox, 2006). However, it is not until the Late PPN-early Late Neolithic (7500-6300 cal BCE) that these morphotypes become dominant at sites across the Fertile Crescent, indicating that this trait had become fixed in cereal populations (Arranz-Otaegui et al., 2016; Tanno and Willcox, 2006).

The southern Levant is an important region for the study of early plant management and domestication in southwest Asia, with archaeobotanical evidence indicating that wild barley (*Hordeum spontaneum*) and wild emmer wheat (*Triticum dicoccoides*) were brought into cultivation and independently domesticated here in the Early Holocene (Arranz-Otaegui et al., 2016). Located 20 km apart in the Wadi el-Hasa in southern Jordan, the archaeological sites of Sharara and el-Hemmeh, provide a significant opportunity to examine how barley and emmer were managed at a local level during the Early Holocene and to track changes in their domestication status (Fig. 1). The PPNA settlement of Sharara (c. 9250-9200 cal BCE) is situated on a rocky slope, high above the wadi floodplain and consists of a loose scatter of structures spread over an area of c. 0.5 ha (Whitlam et al., 2023). el-Hemmeh lies c. 20 km upstream of Sharara on an alluvial fan overlooking the wadi floodplain. The earliest settlement documented at the site dates to the latter half of the PPNA (c. 9400-8700 cal BCE) and is characterised by multiple free-standing and semi-subterranean circular and semi-circular buildings (Makarewicz et al., 2006; Makarewicz and Rose, 2011). A Late PPNB settlement (c. 7500-7000 cal BCE) is also documented and characterised by agglomerated rectilinear architecture (Makarewicz et al., 2006; Makarewicz and Austin, 2006). Analyses of charred plant macroremains from both sites have indicated that inhabitants exploited a range of wild cereals and pulses along with fig (*Ficus carica*) and pistachio (*Pistacia* sp.) (White, 2013; White and Makarewicz, 2012; Whitlam et al., 2023). At Sharara and PPNA el-Hemmeh, barley is the principal cereal crop with glume wheats (primarily emmer) present in low numbers. In contrast, at LPPNB el-Hemmeh, emmer is the major cereal crop with barley a relatively minor component of the assemblage (SI Text, Fig. S1). PDC of barley has previously been inferred at PPNA el-Hemmeh based on the size of grains, the presence of potential arable weeds, and rachis morphology (White and Makarewicz, 2012).

**Figure 1.**
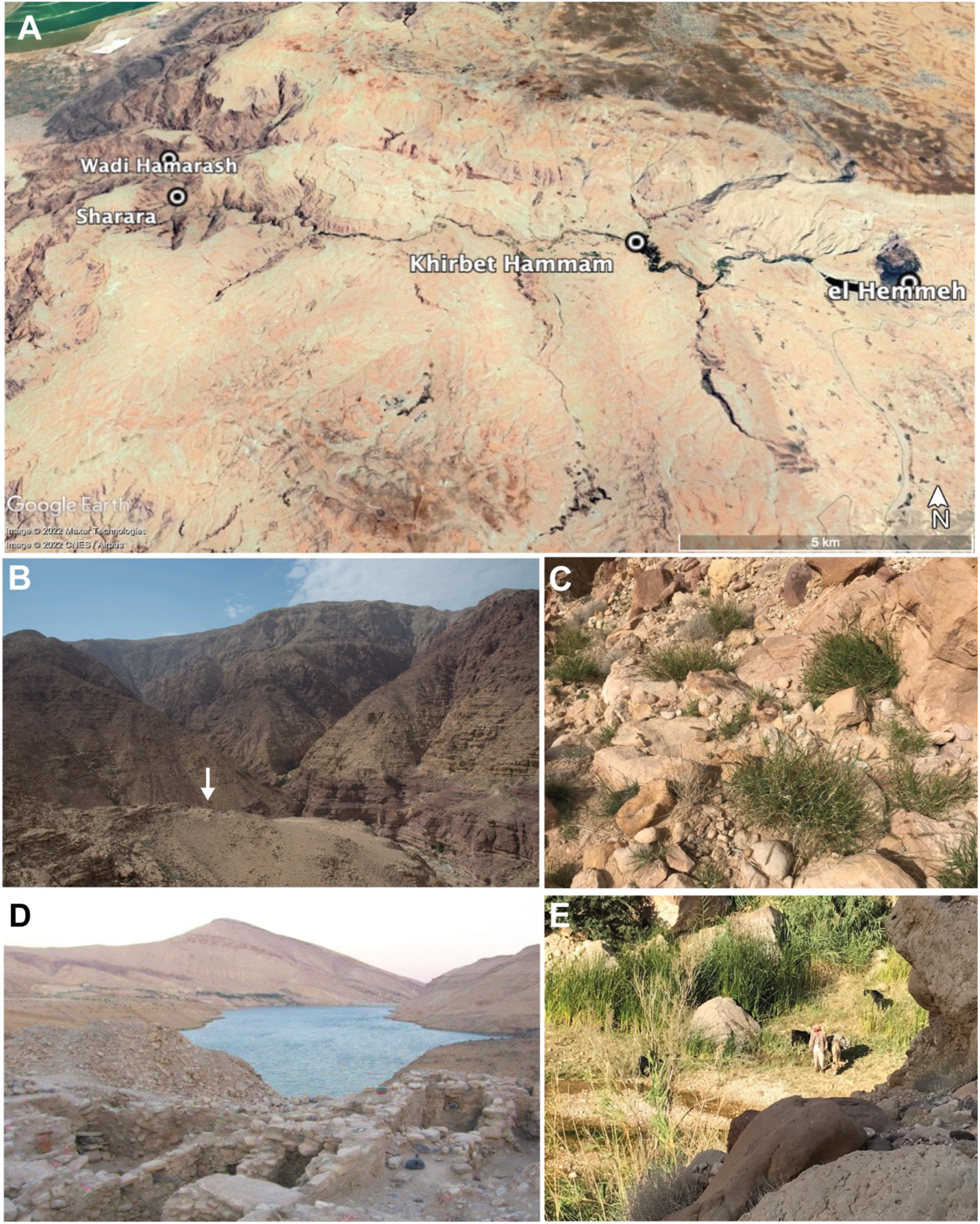
Setting of archaeological sites. (A) Google Earth map showing the location of Sharara and el-Hemmeh in the Wadi el-Hasa. (B) Position of Sharara, indicated by the arrow, in the Wadi el-Hasa. (C) Sparse vegetation growing on rocky slopes around Sharara. (D) Position of el-Hemmeh on an alluvial terrace overlooking the wadi. (E). Rich vegetation growing around fertile wadi edges. [both Sharara]

In this study we report on additional analyses undertaken at both sites. This includes morphological and metric analysis of cereal remains (barley and emmer) to track loss of shattering and increased grain size traits, stable carbon isotope analysis to elucidate water availability, and weed ecological analysis to reconstruct cereal growing conditions.

### Analysis of rachis morphology

To track the domestic status of barley and emmer at Sharara and el-Hemmeh we examined all rachis remains and classified these according to the appearance of their abscission scars. In wild wheat and barley, which have evolved so their ears spontaneously shatter upon maturity, plants produce shattering rachis morphotypes that are identified by their smooth abscission scars (Colledge, 2001, p. 65; Tanno and Willcox, 2012)(Fig. 2a). In domestic wheat and barley, the ears have evolved to remain intact upon ripening, meaning it is necessary to mechanically separate these to free the grain (i.e., by threshing). This results in distinct non-shattering rachis morphotypes that are identified by their rough abscission scars (Colledge, 2001, p. 65; Tanno and Willcox, 2012)(Fig. 2c). For emmer, a secure identification of domestic status is also possible by analysing the longitudinal section of the upper abscission scar. In shattering (wild) emmer this appears flat in orientation, and in non-shattering (domestic) emmer (*Triticum dicoccum*) this appears lifted from the internode (Weide et al., 2015).

**Figure 2.**
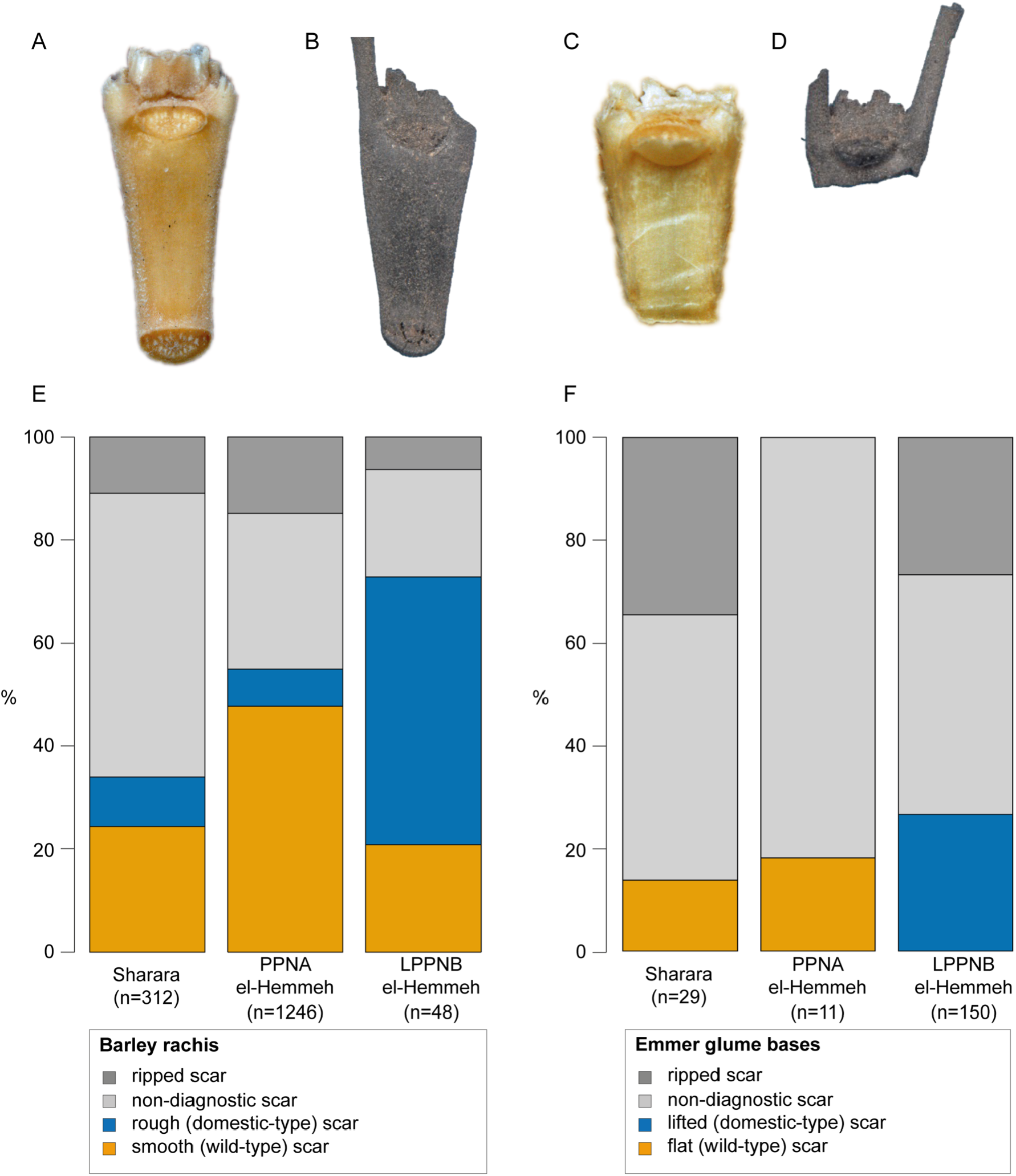
Rachis morphotypes. (A) Modern example of wild barley rachis internode showing smooth abscission scar. (B) Archaeological specimen from el-Hemmeh with smooth (wild-type) abscission scar. (C) Modern example of domestic barley rachis showing rough abscission scar. (D) Archaeological specimen from el-Hemmeh with rough (domestic-type) abscission scar. (E) Relative percentages of different barley rachis morphotypes at Sharara (n=312), PPNA el-Hemmeh and LPPNB el-Hemmeh, with total numbers of specimens shown. (F) Relative percentages of different emmer rachis morphotypes at Sharara, PPNA el-Hemmeh and LPPNB el-Hemmeh with total numbers of specimens shown.

For barley, we distinguished between smooth, rough, non-diagnostic and ripped abscission scars, and for emmer, between flat, lifted, non-diagnostic and ripped abscission scars (see materials and methods). Ripped abscission scars likely result from cereal processing and in terms of assigning domestic status are considered non-diagnostic (SI Text). Figure 2 illustrates the relative proportions of different rachis morphotypes for barley and emmer within each assemblage. These results clearly demonstrate that PPNA barley and emmer are morphologically wild at both sites, with no lifted scars identified from emmer remains, and only low proportions of rough scars identified for barley. The presence of low proportions of non-shattering rachis in wild barley populations is consistent with studies that report the minor occurrence of rough scars (c. 10%) in modern wild barley populations (Kislev, 1989). At LPPNB el-Hemmeh, both barley and emmer appear morphologically domesticated based on the rachis remains. While all diagnostic LPPNB specimens of emmer produced the lifted scar associated with non-shattering, 71% of the diagnostic specimens of barley had a rough abscission scar. This indicates that although the non-shattering trait had yet to become fixed in barley at LPPNB el-Hemmeh, it was present at a far higher proportion than could be maintained in wild populations. This is consistent with evidence from other M/LPPNB sites in the Levant (Arranz-Otaegui et al., 2016).

### Grain size analysis

To investigate variation in barley grain size, we measured breadth and thickness of all grains and grain fragments. We also classified grains into three categories: ‘wild’, ‘intermediate’ and ‘domestic’, based on size parameters previously applied at el-Hemmeh (see Materials and methods; SI Text). The results are presented in Figure 3. Due to the lack of securely identified grains at Sharara and PPNA el-Hemmeh, we did not undertake metrical analysis of emmer.

**Figure 3.**
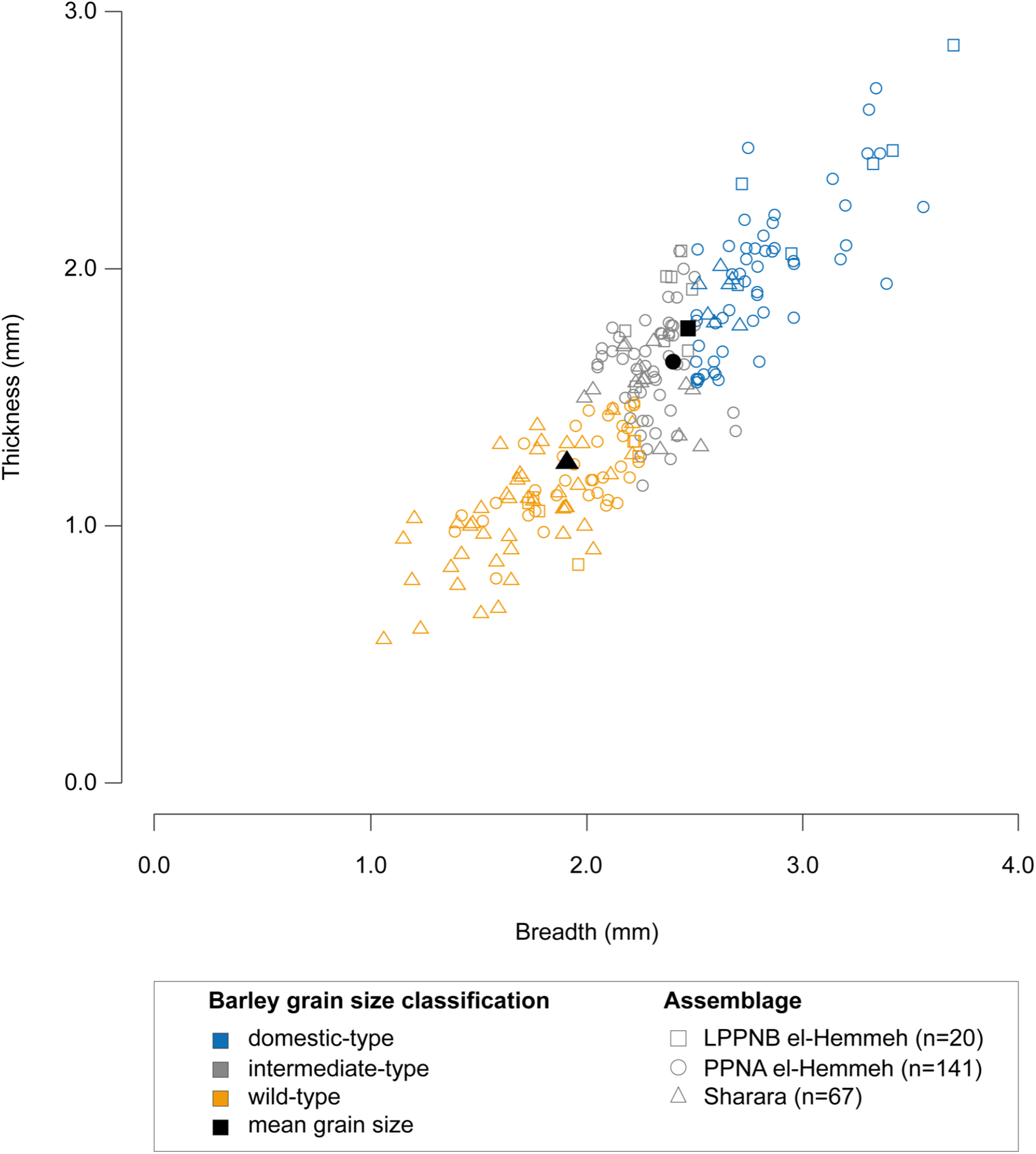
Scatterplot showing the breadth and thickness of 228 barley grains at Sharara, PPNA el-Hemmeh and LPPNB el-Hemmeh, coded according to whether these are classified as ‘wild’, ‘intermediate’ or ‘domestic’ as described in text. Average grain size for each assemblage is indicated by solid black symbols.

Our results demonstrate that the size of barley grains varies both within and between sites, with all three assemblages showing overlap in grain size and producing barley grains classified as wild, intermediate and domestic (Fig. 3). The smallest grains were produced by Sharara, where the average grain size (1.91 x 1.25 mm) sits comfortably in the wild range. At el-Hemmeh LPPNB barley grains (2.47 x 1.77 mm) were larger, on average, than barley grains from PPNA levels (2.40 x 1.64 mm). However, the size difference observed between barley grains at PPNA and LPPNB el-Hemmeh was relatively small compared to the difference observed between barley grains at Sharara and el-Hemmeh. Both PPNA and LPPNB el-Hemmeh also produced a similar proportion of grains that were classified as wild and domestic, despite rachis evidence indicating a shattering (wild) barley population in the PPNA, and a non-shattering (domestic) barley population in the LPPNB. This suggests that differences in grain size at our sites can be better explained by variation within and between populations than by variation resulting from domestication. In other words, phenotypic plasticity and genetic diversity play a greater role in determining grain size than domestication.

### Stable carbon isotope analyses

To explore variation in growing conditions at Sharara and el-Hemmeh, and its potential effect on grain size, we undertook stable carbon isotope analysis of barley and glume wheat grains. This provides a means of inferring the different moisture conditions under which the grains developed. The results of stable carbon isotope analysis are presented in Figure 4, with barley Δ^13^C values presented as single grain values grouped by the wild, intermediate and domestic size categories applied above (see SI Text). Comparing Δ^13^C values of barley grains at el-Hemmeh, the larger, ‘domestic’ grains are higher in Δ^13^C than the wild-type grains, although considerable overlap is present for individual grain values (Fig. 4A). Here, a statistically significant (p<0.001) but weak (r^2^=0.24) correlation is present between the size of the barley grains and Δ^13^C values. Therefore, water availability at el-Hemmeh is correlated to grain size, although not as the only contributing factor. At Sharara such a pattern is not obvious, although the lowest (‘driest’) Δ^13^C values are present for wild grains.

**Figure 4.**
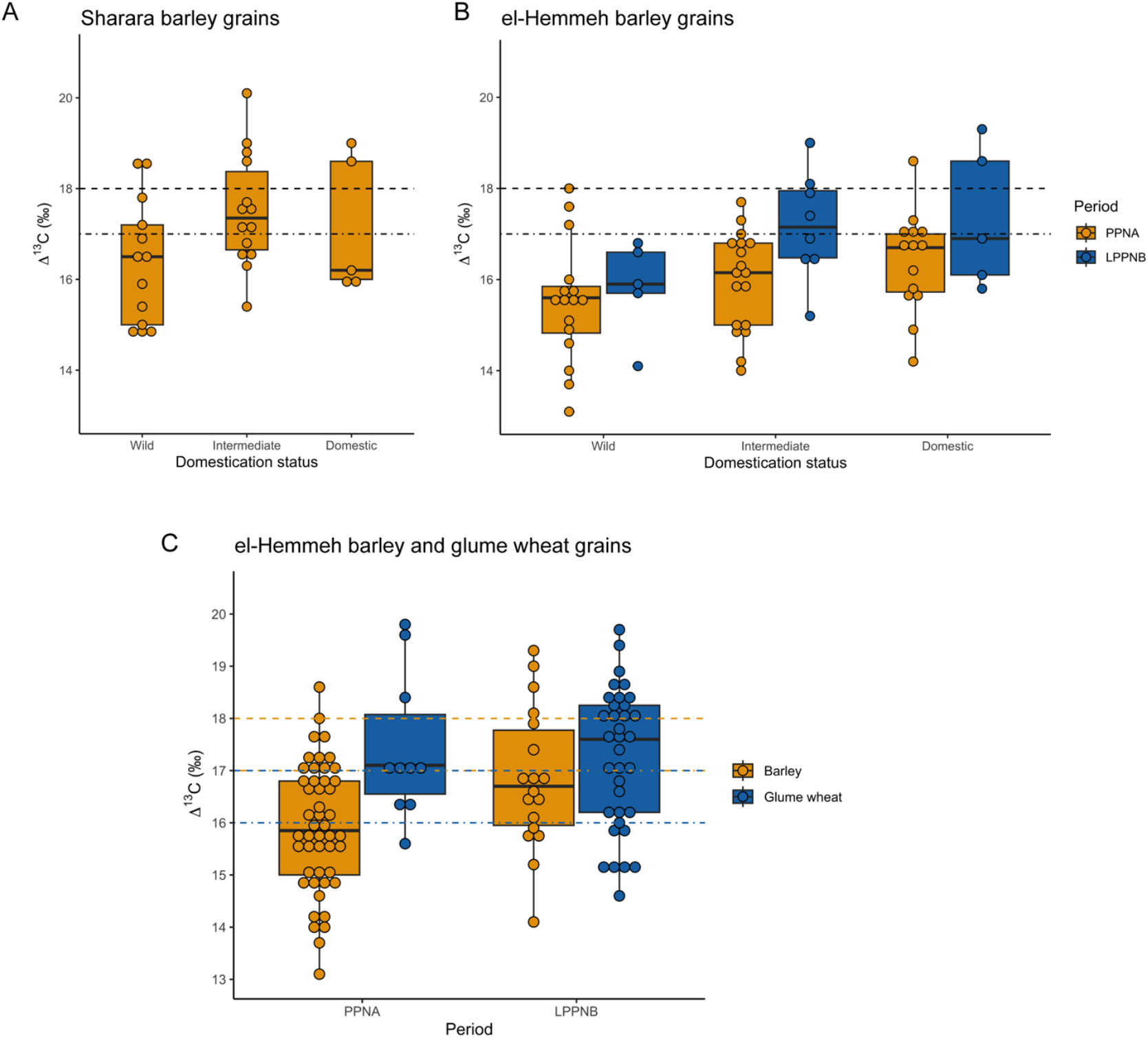
Bar charts of carbon stable isotope discrimination. (A) Bar charts, with circles for individual grain values, showing Δ^13^C in ‰ by domestication status according to White (2013, p. 96) criteria for PPNA barley grains from Sharara. (B) Bar charts with circles with individual grain values, showing Δ^13^C in ‰ by domestication status according to White (2013, p. 96) criteria for PPNA (orange-yellow/light) and LPPNB (blue/dark) barley grains from el-Hemmeh. (C) Bar charts, with circles for individual grain values, showing Δ^13^C in ‰ for barley (orange-yellow/light) and glume wheat (blue/dark) for both the PPNA (left) and the LPPNB (right).

We also observe that barley Δ^13^C values are elevated at (PPNA) Sharara compared to PPNA (but not LPPNB) el-Hemmeh (t-test, p<0.001), indicating that the barley grew under wetter conditions at Sharara, despite the fact that grains are smaller on average, likely in adaptation to local aridity. There are multiple potential reasons, except for Sharara receiving slightly more rainfall because of its location (currently c. 50 mm more annually); the plants could have grown in a wetter location (for example closer to a wadi bed for the Sharara plants and growing higher on slopes at el-Hemmeh); or they could simply have come from a wetter year or a year with more regular rainfall.

Barley Δ^13^C values are elevated at LPPNB el-Hemmeh compared to PPNA el-Hemmeh (t-test, p=0.02) (Fig. 4A), indicating that in the later period more barley plants grew in wetter conditions than during the PPNA. This potential shift in water availability for barley between PPNA and LPPNB el-Hemmeh signals a change in growing conditions that might relate to cultivation. This inference is strengthened by the high proportions of non-shattering (domestic-type) barley (and emmer) rachis in the LPPNB assemblage, which could only be maintained under cultivation. At Sharara and PPNA el-Hemmeh, it is possible that the larger domestic-sized grains may derive from plants that grew in wetter conditions as a result of cultivation, but this may also relate to natural variation in water availability.

Although no clear evidence is present for a shift in water availability in the glume wheat, it is interesting that glume wheat appears ‘wetter’ than barley at least for the PPNA (t-test, p=0.008), and about equally wet for the LPPNB at el-Hemmeh (Fig. 4B). This indicates that glume wheats were growing in different conditions than barley, since under similar conditions (in modern studies) barley tends to have higher Δ^13^C values, as it is more drought-resistant than wheat (Ferrio et al., 2005; Flohr et al., 2019; Wallace et al., 2013).

### Weed ecological analyses

To test whether cultivation, specifically tillage, had an effect on grain size, we undertook functional weed ecological analysis to explore whether cereals were growing in tilled or untilled environments, using the model developed by Weide et al. (2022). This model uses the functional traits of potential arable weeds relating to mechanical soil disturbance, which have been successful in distinguishing between modern-day arable (tilled) and non-arable (untilled) cereal habitats, to locate archaeobotanical samples along an ecological continuum of low to high soil disturbance (SI Text). Figure 5 shows the relationship of modern arable (tilled) fields and non-arable (untilled) wild cereal habitats in Israel to the discriminant function extracted to distinguish between these regimes, using flowering duration as the sole discriminating variable. The discriminant function was then used to classify archaeobotanical samples from Sharara (n=2) and PPNA el-Hemmeh (n=7) that had a high probability of representing crops and their associated weeds, based on the numbers of cereal and weed items they contained. Samples have additionally been coded according to whether they had a high or low likelihood of representing crops and their associated weeds based on crop processing analysis. No samples from LPPNB el-Hemmeh met the criteria for inclusion (SI Text).

**Figure 5.**
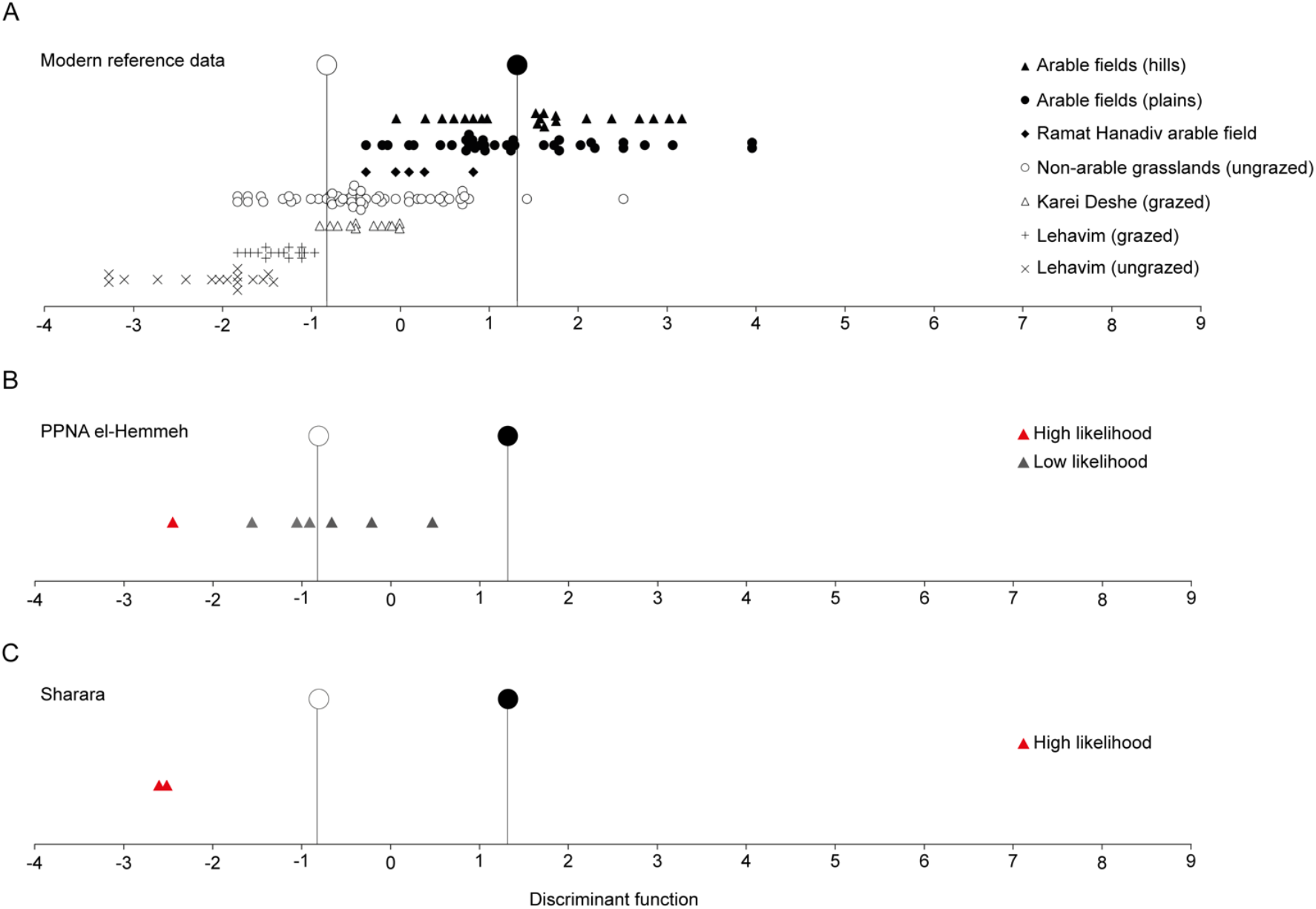
Discriminant analysis to distinguish between arable (tilled) and non-arable (untilled) habitats, with the larger closed and open symbols in each panel indicating the group centroids based on modern data. (A) modern data from arable (closed symbols) and non-arable (open symbols) reported in Weide et al.(2022), the larger closed and open symbols in each panel indicate the group centroids based on modern data. (B) Samples from PPNA el-Hemmeh. (C) Samples from Sharara. Archaeological samples coded according to how likely it is that the sample represents crops and their associated weeds on the basis of crop processing analysis

For Sharara, both samples were classified as non-arable in the DA, indicating that cereals were growing in conditions of low mechanical soil disturbance. Six out of seven PPNA el-Hemmeh samples were also classified as untilled, with only one sample classified as tilled in the DA indicating higher levels of mechanical soil disturbance. However, it is worth noting that the discriminant score for this sample (0.455) overlaps with modern-day non-arable (untilled) cereal habitats (Fig. 5A), and that the sample had a low likelihood of representing crops and their associated weeds on the basis of crop processing analysis. Only one PPNA el-Hemmeh sample was considered to have a high likelihood of representing crops and their associated weeds on the basis of crop processing analysis, and this was classified as untilled in the DA (Fig. 5B). These results indicate that cereals at Sharara and PPNA el-Hemmeh were growing in relatively low disturbance environments and would not have been subject to systematic annual tillage.

## Discussion

Based on the analyses undertaken for this study, we argue that cereal evolution in the southern Levant (as evidenced by loss of shattering and increased grain size) was a product of both genetic selection and developmental plasticity. Our results provide evidence for increased grain size in the PPNA, with larger (domestic-sized) barley grains present at both Sharara and PPNA el-Hemmeh. These larger grains occur alongside shattering rachis morphotypes – a combination that is typically interpreted as evidence for a population under PDC (Colledge et al., 2018; Fuller, 2007; Willcox, 2004). However, weed ecological analysis at Sharara and PPNA el-Hemmeh has demonstrated that cereals were growing in conditions of low disturbance, consistent with those found in modern-day untilled wild cereal stands. We therefore reject the hypothesis that the presence of larger grains at our PPNA sites represents genetic selection for increased grain size under cultivation (i.e. tillage). Instead, we propose that differences in grain size at our sites reflect a plastic response to variation in growing conditions, specifically water availability, since stable carbon isotope analysis indicates a relationship between grain size and moisture. Our results also suggest that while plasticity is important in determining grain size *within* a population, between populations its effect is mediated by genetic variation. This accounts for barley grains at Sharara being smaller on average than those at el-Hemmeh despite growing in wetter conditions. In other words, there is a genetic and plastic component to grain size.

At LPPNB el-Hemmeh, genetic selection for indehiscence is evidenced by the high proportions of non-shattering barley and emmer rachis. This is consistent with evidence from across the Fertile Crescent for the establishment of domestic cereal populations during the Late PPN-early Late Neolithic (Arranz-Otaegui et al., 2016). While it has not been possible to undertake weed ecological analysis at LPPNB el-Hemmeh, the high proportions of non-shattering rachis provide indirect evidence for cultivation, as this trait could only be selected for and maintained with sustained human intervention in the plants’ life cycle. Alongside this evidence for cultivation and domestication, we find an increase in grain size at LPPNB el-Hemmeh from PPNA levels at the site. This could reflect selection for larger grains under cultivation (i.e. genetic selection). However, this increase in grain size also coincides with an increase in wetness at LPPNB el-Hemmeh, as evidenced by stable carbon isotope analysis. An alternative explanation, therefore, would be that cultivation created conditions in which cereal grain size could be maximized due to plasticity.

Taken together, the evidence we have presented here challenges conventional models of plant domestication in southwest Asia, in which increased grain size is selected for prior to the loss of shattering, and where the presence of large (domestic-sized) grains at a site provides evidence for PDC. We argue instead, that increased grain size in the Early Holocene reflects phenotypic plasticity in response to more favourable (specifically wetter) growing conditions. We do not rule out the possibility that human management and modification of the environment helped to create favourable growing conditions. However, during the PPNA this is unlikely to have involved systematic annual tillage and PDC as traditionally formulated, based on our findings. In this scenario, genetic selection is not required to explain increased grain size prior to loss of shattering. We propose instead that genetic selection for increased grain size occurred after (or during) selection for non-shattering rachis, with cultivation and harvesting known to relax selection pressures relating to dispersal, which tend to advantage small seeds that can disperse further (Brown et al., 2009). This is supported by experimental studies, which have suggested that larger-seeded genotypes might have gained a selective advantage only when selection for dispersal was relaxed (i.e. once you have non-shattering)(Preece et al., 2017).

These results open-up the possibility that increased grain size resulting from developmental plasticity may have played a role in directing cereal evolution in southwest Asia, as argued by the EES. Plasticity-led evolution, suggested by an increasing number of studies, argues that the plastic expression of phenotypic variation can contribute to the domestication process by enhancing co-evolutionary relationships, which ultimately lead to the fixation of these traits (Moczek et al., 2011; Ng and Kinjo, 2023; Pfennig et al., 2010; West-Eberhard, 2003). Our study underlines the need for further research, to clarify *how* plastic increases in grain size translated into genetic changes in grain size and the nature of these enhanced co-evolutionary relationships. Here we agree with a number of recent studies that argue for more focused investigations into the complex evolutionary dynamics involved in plant domestication, the constellation of traits this involved and its wider ecological context (Jones et al., 2021; Preece et al., 2021; Zeder, 2018, 2017).

## Materials and Methods

All archaeobotanical samples used in this study (n=80) were analysed at the School of Archaeology, University of Oxford. Charred macrobotanical remains were identified in comparison to modern reference material housed at the School of Archaeology, using a Leica MZ75 stereomicroscope at magnifications of 6x to 50x (SI Text). A single sample from PPNA el-Hemmeh emerged as an outlier in initial data analysis and has been excluded from the analyses presented here (SI Text, Fig. S1-3). Measurements of barley grains were obtained using either a PixeLINK camera attached to a Nikon SMZ25 stereomicroscope and coupled to a PC, assisted by PixeLINK Capture and NIS-Elements D software, or a Leica DFC495 camera attached to a Leica Z6 APO and coupled to a PC, assisted by LAS software. Measurements were used to classify grain as ‘wild’, ‘intermediate’ and ‘domestic’ following size parameters previously applied to barley grain at el-Hemmeh by White (2013, p. 96). We also experimented with classifying barley grains into the same categories based on different parameters applied by Colledge (2001, p. 65) at sites in the southern Levant (SI Text, Fig. S4-5).

Individual grains were analysed for their carbon isotopic composition, 32 for Sharara, mostly from Space 5 (Whitlam et al., 2023) with some additional contexts, 60 from 7 contexts for PPNA el-Hemmeh, and 51 from 9 contexts for LPPNB el-Hemmeh (details in SI). No conclusions are drawn based on individual grains or individual contexts. Each grain was photographed in ventral, lateral, and cross-section views and assessed for their state after charring (Stroud et al., 2023). Subsequently over 10% of the grains were analysed using Fourier Transform Infrared Spectroscopy (FTIR) and their spectra assessed for potential contamination to decide on the appropriate pretreatment (Styring et al., 2016; Vaiglova et al., 2014). Each grain was then pretreated with 0.5M HCl, weighed into tin capsules, and analysed on a Sercon EA-GSL at the University of Oxford’s Research Laboratory for Archaeology and the History of Art. Using international and in-house standards, the results were calibrated. The standard uncertainty at 1sd was ≤0.2‰ (mostly ≤0.1‰). Details of the methods and materials can be found in the SI Text.

For weed ecological analysis, following Weide et al. (2022), flowering duration was selected as the most useful trait for distinguishing between tilled and non-tilled habitats. We considered using vegetative propagation as an additional trait to inform the analysis but this was not possible based on the data and material available (SI Text). Annual and perennial species that flower for an extended period have a high probability of regenerating from seeds in high disturbance regimes, and this trait has been related to conditions of high mechanical disturbance in different climate and ecological settings, including England, Greece, Spain and Morocco, demonstrating that this is functionally independent from local environmental conditions and applicable to archaeological contexts (Bogaard, 2004; Bogaard et al., 2016; Charles et al., 2002; Hamerow et al., 2020; Jones et al., 2000). All samples containing at least 30 cereal remains and 10 weed seeds that were identified to a taxonomic level that allowed attribution of a functional-trait value were included in the analysis. We also experimented with using a minimum of 100 cereal items as a cut-off for inclusion in the analysis (SI Text; Tables S1-3). The process of assigning flowering duration values to taxa was done using information from the *Flora Palaestina* and is detailed in the SI Text (Tables S4-5). For taxa that weren’t identified to species, an average flowering duration value was calculated based on a list of possible species identifications. Where flowering duration varied by more than two months between possible species, we also tested the minimum and maximum flowering durations in the analysis (Fig. S8). The results of crop-processing analysis to determine how reliable samples were in terms of their crop-weed relationships is reported in the SI (SI Text, Fig. S6-7, Tables S2-3).

Note on data, materials and software availability: Data processing was done in MS Excel and R. Data analysis was undertaken in R Studio versions 4.1.2, 2023.12.1, and 2024.04.2+764. For weed ecological analysis, data were prepared in R studio with the discriminant analysis performed using IBM SPSS version 27. All archaeobotanical samples received a discriminant score, which we used to visualize the relative position of samples to each other by plotting the samples along the extracted discriminant function (in Microsoft Excel). Average attribute scores for each analysed modern and archaeobotanical sample and the outputs of the DAs are provided in Table S7. Figures in the main text have been edited using Adobe Illustrator. The underlying data as CSV and XLSX files, an accompanying README file for each dataset, and an R Script to replicate all graphs and tables in the main text and SI are provided at the University of Oxford data repository.

## Acknowledgments

We thank all members of the Sharara and el-Hemmeh team for their help in collecting samples, particularly Dr. Chantel White (University of Pennsylvania) for her work at el-Hemmeh. We also thank Elizabeth Stroud and Peter Ditchfield (University of Oxford) for their advice on the FTIR and stable isotope analyses and interpretation. This research was funded by a British Academy Postdoctoral Fellowship (pf170053, J.W.), the National Geographic Council for Exploration and Research (HJ-043R-17, C.M.) and the Deutsche Forschungsgemeinschaft (EXC 2150, ROOTS – Social, Environmental, and Cultural Connectivity in Past Societies; University of Kiel). Permission to excavate at Sharara and el-Hemmeh was granted by the Department of Antiquities of Jordan, who also authorised the export of archaeobotanical samples (Jordanian Department of Antiquities Export Permit No. 12/5/4056 and 12/5/2180).

I have been occasionally in touch with my teammates, I’ve been really lucky that many of them have reached out to me over the years to say ‘hello’ and check in to how things are going. Between living overseas and regular fieldwork that takes me to remote places where only a satellite phone is the only means of communication, it can be tough to remain linked in with folks back in the US. But the now archaic medium of Facebook has been the primary way we’ve been able to connect, and it’s been wonderful seeing their relationships bloom, their families grow, and their successes take flight.

