## Supplemental Information for "Developmental plasticity and genetic selection shaped cereal evolution in the Early Holocene southern Levant"

*(Pre-print version for discussion)

Jade Whitlam[1,2*], Pascal Flohr[3,1], Amy Bogaard[1,4], Mike Charles[1], Bill Finlayson[1] and Cheryl A. Makarewicz[3*]

[1]School of Archaeology, University of Oxford, 1 South Parks Rd, OX1 3TG, Oxford, United Kingdom

[2]Department for Continuing Education, University of Oxford, OX1 2JA, Oxford, United Kingdom

[3]Institute for Prehistoric and Protohistoric Archaeology, University of Kiel, Johanna-Mestorf Strasse 2-6, D-24118, Germany

[4]Sante Fe Institute, Hyde Park Road, Sante Fe, NM 87501, USA

**Corresponding author:**

Jade Whitlam

Cheryl Makarewicz

**Supporting materials for this manuscript include the following:**

Supporting text
Figures S1 to S10
Tables S1 to S11
Legends for Datasets S1 to S7
SI References

Datasets S1 to S6: available at the University of Oxford data repository

Dataset S7: available at the University of Oxford data repository

### Supporting Information Text

**Part 1** (Section author: Jade Whitlam)

#### Sampling and recovery of charred macrobotanical remains

Charred plant remains were recovered at Sharara and el-Hemmeh through a programme of systemic sampling and flotation. At el-Hemmeh, up to 10L sediments recovered from midden, hearth and storage contexts were floated using a hand-pump system that gently agitated sediments using continuously replaced water drawn from the reservoir adjacent to the site (Shelton and White, 2010). Floated materials were recovered using a 250µm mesh. Initial analysis of the charred macrobotanical remains recovered from el-Hemmeh during the 2004-2007 seasons, are reported in White and Makarewicz (2012) for PPNA plant remains and White and Wolff (2012) for LPPNB remains. For this study, we focused on a subset of previously studied PPNA (n=8) and LPPNB (n=11) samples, which were re-analysed to ensure consistency in identification and quantification between assemblages, using the methods reported in Whitlam et al. (2023). We also analysed 27 previously unstudied samples from PPNA levels at el-Hemmeh collected between 2010-2014. At Sharara, up to 10L of sediments were floated in buckets making use of the perennial water flow in the Wadi el-Hasa. All Sharara samples (n=34) analysed by Whitlam et al. (2023) were included in this study. Sorting and identification of charred plant remains took place in in the Archaeobotany Laboratory at the School of Archaeology, University of Oxford.

#### Identification and documentation

All non-woody charred plant remains were identified using a Leica MZ75 stereomicroscope at magnifications of 6× to 50×. Photographs and measurements of charred plant remains were obtained at the School of Archaeology using either a PixeLINK camera attached to a Nikon SMZ25 stereomicroscope and coupled to a PC, assisted by PixeLINK Capture and NIS-Elements D software, or a Leica DFC495 camera attached to a Leica Z6 APO and coupled to a PC, assisted by LAS software. Identifications were made according to morphological characteristics, surface texture and size, and verified by comparison to modern reference material housed at the School of Archaeology. Plant remains were quantified using the principle of recording the 'minimum number of individuals' (MNI) following Jones (1991). Nomenclature follows the *Flora Palaestina* (Feinbrun-Dothan, 1986, 1978; Zohary, 1972, 1966), except for family names where modern conventions have been applied. Full sample-by-taxa data are provided in Datasets S1 and S2.

#### Archaeobotanical overview

To explore the botanical composition of samples, stacked bar charts showing the relative proportions of major botanical categories within samples containing 10 or more items were produced for Sharara, as well as PPNA and LPPNB el-Hemmeh (Fig. S1). These clearly demonstrate that barley was the major cereal taxon at both Sharara and PPNA el-Hemmeh (Fig. S1A, C), while at LPPNB el-Hemmeh (Fig. S1E) this was glume wheat. At PPNA el-Hemmeh, sample HEM_1143 (the mixed midden and ash dump fill of Cut 1142), stands out for its high proportion of glume wheat and relatively low proportion of barley.

There are also differences in the parts of barley represented at Sharara and PPNA el-Hemmeh, with barley rachis dominating at Sharara and barley grain dominating at PPNA el-Hemmeh (Fig. S1B, D). Only three samples at PPNA el-Hemmeh (HEM_3364, HEM_3476A and HEM_3476B) produced high proportions of barley rachis. At LPPNB el-Hemmeh the majority of the glume wheat consisted of glume bases rather than glume wheat grains (Fig. S1F). This was also true of the glume wheat-rich sample HEM_1143 from PPNA levels (Fig. S1D).

#### Correspondence analysis

We undertook correspondence analysis to further explore similarities and differences in the botanical composition of samples from PPNA and LPPNB el-Hemmeh. A list of samples and whether these represent PPNA or LPPNB deposits is provided in Dataset S3. We were particularly interested to see how HEM_1143 behaved in this multivariate analysis on the basis of all taxa, rather than the simplified botanical categories used to draw comparisons in Fig. S1. All taxa occurring in five or more (i.e. 10%) of el-Hemmeh samples (n=38), and all samples containing 30 or more items representing these taxa (n=31) were included in the correspondence analysis. The sample plot is presented in Fig. S2A. This clearly demonstrates that sample HEM_1143 groups with LPPNB samples to the bottom right of the plot, reflecting the association of these samples with glume wheats and small-seeded legumes, which are pulled in this direction in the corresponding species plot (Fig. S2B). In contrast, to sample HEM_1143, the majority of PPNA samples are distributed along the positive (top) end of the second axis and close to the origin of axis 1. This reflects their association with a combination of barley grain, pulses and various fruit/nut taxa. Three PPNA samples (HEM_3364, HEM_3476A and HEM_3476B) diverge from this group and are in the lower left quadrant, reflecting their strong association with barley rachis as previously noted. This initial data analysis indicates that sample HEM_1143 is more similar to LPPNB samples in terms of its botanical composition than PPNA samples. Although context 1143 itself is securely PPNA, given this uncertainty, and in the absence of direct radiocarbon dates supporting the PPNA origin of this material, sample HEM_1143 has been excluded from the final analyses.

**Analysis of chaff morphology**

For this study, *Hordeum spontaneum/vulgare* (barley) rachis internodes were identified as exhibiting either a smooth (wild-type) or rough (domestic-type) abscission scar, following criteria described by Colledge (2001, p. 65) and Tanno and Willcox (2012). If the abscission scar could not be confidently identified as belonging to one of these types, usually due to poor preservation of the scar tissues, the scar was recorded as ‘non-diagnostic’. An additional ‘ripped’ rachis category was used to record specimens that were missing part, or all, of their abscission scar. We interpret this as evidence of pounding to dehusk grain (Tanno and Willcox, 2012; Whitlam et al., 2023). These ripped rachis remains were considered non-diagnostic in terms of their domestic status. Glume bases of *Triticum dicoccoides/dicoccum* (emmer wheat) were identified as exhibiting either a flat (wild-type), lifted (domestic-type) or non-diagnostic abscission scar, following criteria laid out by Weide et al. (2015). Ripped specimens, which displayed a morphology similar to that described for barley above, were also recorded and are considered as non-diagnostic in term of their domestic status.

The relative percentages of different chaff types for barley and emmer were then calculated for each assemblage (not including sample HEM_1143) (Fig. S3A, B). It is worth noting, that the inclusion of glume wheat-rich HEM_1143 did not alter the pattern observed for barley rachis, as it produced few barley remains. However, it had a significant effect on the pattern observed for emmer at PPNA el-Hemmeh (Fig. S3C, D). This is because HEM_1143 produced the majority of glume bases found in PPNA samples, with a high proportion of these exhibiting the lifted (domestic-type) scar morphology.

**Metrical analysis of barley grains**

Breadth and thickness measurements were taken for all whole barley grains from Sharara, PPNA el-Hemmeh and LPPNB el-Hemmeh. At Sharara, where a relatively small number of whole barley grains were recovered, breadth and thickness were measured for all grain fragments to provide more data points. For PPNA and LPPNB el-Hemmeh, grain fragments were only measured if they were being submitted for stable isotope analysis (carbon and nitrogen). Measurements were taken at the widest point of the grain and are reported in millimeters (mm). Breadth and thickness measurements were used to assign barley grains to ‘wild’, ‘intermediate’ and ‘domestic’ size categories, following parameters previously used by White (2013, p. 96) at el-Hemmeh. Given the somewhat ambiguous nature of these parameters, we repeated this process with parameters previously used by Colledge (2001, p. 64) at other early Holocene southern Levantine sites. In the White classification, barley grains were categorised as ‘wild’ if they had a breadth of 2.25 mm or less and a thickness of 1.5 mm or less, and ‘domestic’ if they had a breadth of 2.5 mm or more and a thickness of 1.5 mm or more. Grains falling between these two groups were categorised as ‘intermediate’. In the Colledge classification, barley grains were categorised as ‘wild’ if they had a breadth of 2.2 mm or less and a thickness of 1.4 mm or less, and 'domestic' if they had a breadth of 2.2 mm or more and a thickness of 1.0 mm or more. Grains falling between these two groups were categorised as 'intermediate'. Results for individual assemblages are shown in Figure S4 and for all grains measured in Figure S5. All measurements of barley grains and their classifications according to White (2013, p. 96) and Colledge (2001, p. 64) are provided in Dataset S4.

**Weed ecology**

Following Weide et al. (2022), flowering duration was selected as the most useful trait for distinguishing between tilled and non-tilled habitats. The use of flowering duration as the sole criterion to distinguish arable from wild-cereal habitats in an archaeological context has been criticised (Willcox, 2023). In their original model Weide et al. (2022) also ran a discriminant analysis (DA) using vegetative propagation along with flowering duration as traits to distinguish between tilled and non-tilled habitats. The inclusion of this second trait resulted in the DA correctly reclassifying 90.2% of all modern samples, compared to 86.7% when only flowering duration was used. While we recognise that a functional ecological model based solely on flowering duration is less powerful than one based on multiple traits, vegetative propagation data was not available for all species considered in this study and would have significantly limited the number of species and samples that were included in the DA.

To ensure that samples included in the weed ecological analysis had a high probability of representing cereal crops and their associated weed assemblages, we only included samples in weed ecological analysis if they contained a minimum of 30 cereal items, and 10 weed items identified to a taxonomic level allowing attribution of a functional trait value (see below). This excluded samples with minor proportions of crop items that are less likely to reflect cereal processing activities and related weed assemblages. We also undertook crop processing analysis to determine whether the cereal and weed remains within each sample were consistent in terms of the stage of the crop processing sequence represented. If consistent samples were considered to have a high likelihood of representing crops and their associated weeds. If inconsistent, the implication is that some weeds may have arrived on site via a route other than cereal harvesting, and thus would not be appropriate for reconstructing the conditions in which cereal crops were growing. Such samples were considered to have a low likelihood of representing crops and their associated weeds (see Table S1).

We first experimented with using a minimum threshold of 100 cereal items and 10 weed items following Weide et al. (2022) for the inclusion of samples in weed ecological analysis. However, lowering the threshold to 30 cereal items did not alter the number of samples included in the analysis for Sharara, while at PPNA el-Hemmeh only four more samples met this lower threshold for inclusion, all having a low likelihood of representing crops and their associated weeds based on crop processing analysis. For LPPNB el-Hemmeh, even with the lower threshold of 30 cereal items, no samples met the minimum criteria for inclusion in weed ecological analysis or crop processing analysis. This was a result of the low numbers of cereal items and the fact that few weed taxa and types could be identified to a taxonomic level sufficient for attribution of a functional-trait value.

**Crop processing analysis.** To determine which stage of the crop processing sequence was represented within archaeobotanical samples, these were compared to ethnobotanical samples from the Greek Island of Amorgos that derive from known stages of the crop processing sequence, in two ways: firstly, in terms of the relative percentages of grain, rachis and weed seeds within samples following Jones (1990), and secondly via weed-based DA following Jones (Jones, 1987, 1984). Samples with minor proportions of crop items (either <100 or <30 depending on the threshold we were using) were excluded from the analyses. As barley and glume wheats have slightly different processing requirements and are unlikely to have been processed together, we also only included samples that represented a relatively pure crop (i.e., that were comprised of at least 80% barley or glume wheat).

For relative percentages of grain, rachis and weed items, results are shown on triangular plots (triplots) alongside the Amorgos samples. Triplots were plotted in R using the ‘CropPro-package’ developed by Stroud et al. (in review). For weed-based DA, remains of wild plant taxa were classified according to their physical characteristics that determine at which stage in the crop processing sequence they are removed, for example, their size, headedness (i.e. the tendency for seeds to stay in heads despite threshing) and aerodynamic properties. These classifications are listed in Datasets S1 and S2 (‘crop pro code’). The size of seeds was taken from measurements based on archaeological specimens and information on seed characteristics collated from various sources, including the *Flora Palaestina*, the Kew seed database (http://data.kew.org/sid/) and other published studies. A discriminant analysis was then performed on both the archaeological dataset and the ethnographic Amorgos samples from known crop processing stages (winnowing by product, coarse sieve by-product, fine sieve by-product and fine-sieve product). This classified each of the archaeological samples into one of the four ethnobotanical groups based on similarities in their weed composition. Only samples with at least 10 weed items that were classified in terms of their physical properties as these relate to crop processing were included in the DA. The DA was carried out in R using the ‘CropPro-package’ developed by Stroud et al. (in review).

**Sharara crop processing analyses.** For Sharara two samples met the threshold of 100 cereal items (SHAR_003 and SHAR_069), with an additional sample (SHAR_077) meeting the lower threshold of 30 cereal items per sample. Considering the relative percentages of grain, rachis and weed items, all three samples appear most closely associated with the Amorgos samples that represent winnowing by-products and coarse-sieve by-products (Fig. S6A). However, only samples SHAR_003 and SHAR_069 produced enough weed items to be included in the weed-based DA. These were both classified as coarse-sieve by-products with a high probability, in agreement with their classification in the triplot (Fig. S6B), meaning both samples were consistent in their crop-weed relationships, and there was a high likelihood these samples represented crops and their associated weeds that were harvested together (Table S2)

**PPNA el-Hemmeh crop processing analyses**

For PPNA el-Hemmeh, five samples met the threshold for inclusion of 100 cereal items (HEM_3237, HEM_3364, HEM_3476A, HEM_3476B, HEM_720). Considering the relative percentages of grain, rachis and weed items, three of the samples (HEM_3364, HEM_3476A, HEM_3476B) appear most closely associated with the Amorgos samples that represent winnowing by-products and coarse-sieve by-products (Fig. S7A). In weed-based discriminant analysis, HEM_3476A was classified as winnowing by-product with a high probability in agreement with its classification in the triplot and is therefore considered to have a high likelihood of representing crops and their associated weeds (Table S3). HEM_3476B was classified as coarse-sieving by-product, in agreement with its classification based in the triplot, but with low probability, while HEM_3364 did not produce enough weed items to be included in the weed-based discriminant analysis. Both samples are, therefore, considered to have a low likelihood of representing crops and their associated weeds (Table S3). The remaining two samples with 100 or more crop items (HEM_3237, HEM_720) did not correspond to any of the ethnographic samples in the triplot, suggesting that they may contain weed items from other sources (Fig. S7A). Both are considered to have a low likelihood of representing crops and their associated weeds. In weed-based discriminant analysis both samples were classified as coarse-sieving by-product with a high probability (Fig. S7B).

When the threshold for inclusion of samples was lowered to 30 cereal items, an additional four samples (HEM_815, HEM_886, HEM_717b, HEM_721A) could be included. All of which are considered to have a low likelihood of representing crops and their associated weeds (Table S3). Three of the samples did not correspond to any of the ethnographic Amorgos samples in the triplot, suggesting that they contained weed items from other sources (Fig. S7A), although in weed-based discriminant analysis these were classified as coarse-sieving by-product with a high probability (Fig. S7B). One sample HEM_717b lay close to the position of Amorgos samples that represent coarse-sieve by-products in the triplot but did not contain enough weed items to be included in weed-based discriminant analysis.

**Reviewing weed taxa for inclusion in weed ecological analysis.** Wild/weed taxa were reviewed prior to weed ecological analysis. All taxa identified to a species-level were included, with information on flowering duration taken directly from the *Flora Palaestina*. For taxa identified to a genus level, potential species lists were drawn up using the *Flora Palaestina*. Any species that could be ruled out on morphological grounds or that were reported as an introduced species in the *Flora Palaestina* were excluded from this list. The remaining species were then input into the analysis as an aggregate species group, using their average flowering duration. If the flowering durations of potential species listed for a taxon varied by more than two months, then the taxon was excluded from the analysis if it was poorly represented at the site in terms of abundance and/or ubiquity. If well represented, however, the taxon was included in the analysis using the average flowering duration of the aggregated species group, after first testing the effect of using the minimum and maximum flowering durations of the aggregated species group in the analysis. Tables S4 and S5 list all the wild/weed taxa identified at Sharara and el-Hemmeh respectively, detailing how these taxa have been treated within the weed ecological analysis conducted for this study. Relevant information from the *Flora Palaestina* is published in Dataset S5.

**Weed ecological analysis**. We used discriminant analysis to distinguish between arable (tilled) and non-arable (untilled) cereal habitats according to their disturbance conditions using the model developed by Weide et al. (2022). The archaeobotanical samples were entered into the classification phase of the analysis as cases with unknown disturbance conditions. The analysis classified each archaeobotanical sample as ‘arable’ (tilled) or ‘non-arable’ (untilled) with a low (<90%) or high (>90%) probability. All archaeobotanical samples received a discriminant score, which was used to visualize the relative position of samples to each other by plotting these along the extracted discriminant function (Fig. S8). Average attribute scores for each analysed archaeobotanical sample and the outputs of the DAs are given in Dataset S6.

Samples from PPNA el-Hemmeh were analysed separately using average, minimum and maximum flowering duration values for aggregate species groups (as outlined above) where differences in flowering duration varied significantly (Fig. S8B). However, while this made some difference to the position of samples on the plot, with maximum flowering duration values pulling samples towards the arable (tilled) centroid, and minimum lowering duration values pulling samples towards the non-arable (untilled) centroid, this difference was marginal. It is clear that whichever values were used for these aggregate species groups, the samples remained distributed towards the untilled end of the spectrum. It should also be noted that the only sample with a high likelihood of representing crops and their associated weeds based on crop processing analysis (HEM_3476A) remained in the same position along the negative (untilled) end of the axis.

**Software and analysis**

For the above analyses all data processing was done in MS Excel and R. Data analysis was undertaken in R Studio version 2023.12.1. For weed ecological analysis, data were prepared in MS Excel and R studio with the discriminant analysis performed using IBM SPSS version 27. An R Script to replicate all graphs and tables in the main text and SI is provided in a data archive [available at the Oxford University Research Archive; DOI archaeobotany dataset forthcoming:].

**Part 2** (Section author: Pascal Flohr)

**Stable isotope analyses**

Carbon and nitrogen stable isotope analysis were conducted on a selection of the grains. Carbon stable isotope analysis can give an indication of the (broad level of) water status of the plant which the grain has derived from (Flohr et al., 2019; Wallace et al., 2013), while nitrogen stable isotope analysis can indicate if the crops were manured (Bogaard et al., 2007; Fraser et al., 2011). While we did not gain $\delta^{15}N$ values for enough grains for the results to be useful for this current study, we are reporting on the methods and average results below nonetheless, and have published the results in a data archive [Oxford University Research Archive; DOI isotope dataset for coming] in case the results can be of use in future research.

**Materials and methods**

**Grain selection:** Grains were selected based on the number available per context (*locus*), or group of contexts, on their condition (completeness and their charring / preservation status), and on minimum weight.

Firstly, the aim was to have a minimum of five grains per context (Flohr et al., 2019; Riehl et al., 2008), and therefore rich contexts were targeted. Nonetheless, to increase the total number of grains per site and period, we also added contexts for which fewer grains were available. We are, however, not using the individual grain values to draw conclusions, and recommend that anyone reusing our results takes the same care.

Secondly, the charring / preservation status of the grains was assessed. Each grain was photographed in ventral, dorsal, and lateral cross-section on either a Leica Z6Apo Z-stacked or on a Nikon (not z-stacked) microscope. The cross-sections were compared to specimens experimentally charred by E. Stroud pers. comm. The condition of each grain was then recorded on a scale of good / borderline / poor (Dataset S7). The aim was to include whole grains in a good condition; however, it was necessary to include borderline and poor grains as well as partial grains. When selecting partial grains, it was made sure that these represented unique individuals. The fact that partial and poor-quality grains were included was taken into account during analysis, since grains categorized as poor potentially provide less reliable isotope values and certain grain parts may also differ in isotopic composition from other parts. Photographs of all grains submitted for stable isotope analysis are available in a data archive [Oxford University Research Archive; DOI grain photo dataset forthcoming].

Finally, the grains were weighed prior to pretreatment and when lighter than 1.4 mg the grains were not used, as it was likely not enough material would be left after pretreatment and further processing. This does mean, however, that we do not have isotope results for the very smallest of the grains.

**Sample size:** An overview of all samples processed for isotopic analysis is available in Dataset S8 and [Oxford University Research Archive, DOI isotope dataset forthcoming: repository dataset Table 1 ]. For Sharara we gained results for 32 barley grains from 10 contexts; 13 of the grains were of the wild type, 14 intermediate type, 5 domestic type following parameters laid out by White (2013). Most of these grains came from Space 5 (n=24) (see (Whitlam et al., 2023)). For el-Hemmeh, results are available for 111 grains, 60 from 7 contexts for the PPNA, of which 48 barley (16 wild, 18 intermediate, 14 domestic) and 10 glume wheat; and 51 from 9 contexts for the LPPNB, of which 18 barley (5 wild, 8 intermediate, 5 domestic) and 35 glume wheat (Table S6). As mentioned above, while the aim was to have at least five grains per context (Flohr et al., 2019; Riehl et al., 2008), due to the low numbers, other available additional contexts were also sampled – however, no conclusions are drawn based on isotope values of individual grains or individual contexts with low numbers of analysed grains.

**Pretreatment and sample preparation**

**Physical cleaning:** The grains were examined at x10-x60 magnification for visible surface contaminants, and a few of the Sharara grains were cleaned to remove adhering sediment [Oxford University Research Archive, DOI isotope dataset forthcoming: Table 1]. This was done by scrapping the dirt off with a razor blade. If possible, cleaning was avoided given the low weight of grains. There were no instances of sediment being found inside the grains. No grains from el-Hemmeh required cleaning.

**Establishing which pre-treatment method to use:** Charred grains may be treated with different pretreatment methods to remove exogenous contaminants from the burial environment, as these can potentially impact the isotope ratios. During the $^{14}$C-dating process, for example, grains and other plant samples are treated with an acid to remove carbonates, an alkali/base to remove humic (and other) acids), and acid again to neutralise the sample (Mook and Streurman, 1983). However, especially the base step removes a lot of endogeneous sample material, which could affect the resulting isotope value, so if not necessary this step is best avoided for stable isotope analysis. To decide on the appropriate pretreatment method, >10% of the grains (11 for el-Hemmeh and five for Sharara) were analysed using Fourier Transform Infrared Spectroscopy (FITR) (see (Vaiglova et al., 2014)). The analyses were conducted on ground samples with an Agilent Carry 640 FTIR instrument with a GladiATR accessory at the Research Laboratory for Archaeology and the History of Art at the University of Oxford. After scanning a sample, using the accompanying Agilent software, the background noise was subtracted, and a baseline correction conducted [Oxford University Research Archive, DOI isotope dataset forthcoming]. Subsequently, the absorbance spectra of the archaeological samples were analysed for potential exogenous contamination by checking if peaks expected for carbonate, nitrate, humic acid, and sulphate were present. Specifically, it was checked if peaks occurred at wavelengths of 720 and 870 cm$^{-1}$ (carbonate), 1085, 1450, and 3300 cm-1 (nitrate), 1010, 1080, and 3690 cm$^{-1}$ (humic acid), and 650 cm$^{-1}$ and 1100 cm$^{-1}$ (sulphate) (Vahur et al., 2016; Vaiglova et al., 2014 ; E. Stroud pers. comm.). None of the samples showed peaks expected for nitrates and sulphates (Fig. S9). Slight carbonate peaks were present in several of the samples at both Sharara and el-Hemmeh. Humic acid peaks at 1010 cm$^{-1}$ and 1080 cm$^{-1}$ were present in some of the samples, but never at 3690 cm$^{-1}$. This is important, since Styring et al. (2016) showed that peaks at 1010 and 1080 cm$^{-1}$ can be caused also by endogenous humics (i.e. from the decomposing grain itself, so with the same isotopic composition, arguably) while a peak at 3690 cm$^{-1}$ was pointing to exogeneous humic contamination (also E. Stroud pers. comm We therefore concluded that the only exogenous contamination present in (some of) the grains were carbonates and that an acid-only treatment to remove carbonates should be sufficient. Moreover, while one could argue to do a full acid-base-acid treatment ‘to be sure’, this would also remove the identified endogenous humics, i.e. a part of the sample.

**Sample preparation:** Following the University of Oxford protocol, each (partial) grain was soaked in 0.5M HCl at 70°C for 40-60 minutes, until the effervescing reaction ceased. Subsequently the samples were rinsed until neutral with ultra-pure water, and then frozen and freeze-dried. The process was started off with whole grains / complete grain parts, but most grains fell apart immediately upon contact with the solvent. When needed, after freeze-drying the samples were ground with an agate mortar and pestle and weighed into tin capsules. 0.64-0.80 mg for ‘carbon-only’ analyses; 0.98-1.10 mg for combined carbon and nitrogen mass spectrometer runs; and up to 5 mg for nitrogen-only analyses for those samples where the nitrogen content of the sample was very low. The aim was for the tin capsule to contain the correct range of carbon and/or nitrogen for the mass spectrometer settings (around 400 µg for carbon and 150 µg for nitrogen).

**Mass spectrometer analysis**

The samples were analysed on a Sercon EA-GSL at the University of Oxford’s Research Laboratory for Archaeology and the History of Art. The international standards EMA-Spruce and EMA-Sorghum were used as standard reference materials for $\delta^{13}$C, EMA-P2 and Leucine for $\delta^{15}$N, and in-house cow and seal collagen as well as alanine for both $\delta^{13}$C and $\delta^{15}$N (Table S7). The alanine standard was dispersed in pairs throughout the run. USGS-40 and USGS-41 standards were also added, but since these yielded clearly erroneous results (large offsets from the other standards) and large standard deviations, they were, after careful deliberation, further discarded. 'Raw' results of each of the runs are available at [Oxford University Research Archive, DOI isotope dataset forthcoming].

**Calibration and corrections**

Following guidelines of the archaeological stable isotope laboratories at the University of Reading (G. Müldner), the University of Oxford, and Szpak et al. (2017), two- and three-point calibrations to the VPDB and AIR scale were conducted, using two to three of the standards. This correction was subsequently checked against the remaining standards. The standard uncertainty for both $\delta^{13}C$ and $\delta^{15}N$ was $\leq 0.2‰$ (mostly $<0.1‰$) at 1 sd. Because one nitrogen-only run showed a very large variation in the alanine standard values, its results were discarded. A linearity correction of 0.13‰ per 100 microgram was applied for the $\delta^{13}C$ values (all original values are available at [Oxford University Research Archive, DOI isotope dataset forthcoming]. Due to the failed $\delta^{15}N$ run and low amounts of available material for $\delta^{15}N$ analysis (which with the Sercon EA-GSL mass spectrometer and its settings required more materials than $\delta^{13}C$ analysis), no such correction could be done for the $\delta^{15}N$ values. Finally, the charring effect was corrected for by subtracting 0.11‰ for $\delta^{13}C$ and 0.3‰ for $\delta^{15}N$ (Nitsch et al., 2015). The processed results of each of the runs are available at [Oxford University Research Archive, DOI isotope dataset forthcoming]. and an overview of all results and their corrected values can be found there as well in Dataset S7.

**$\Delta^{13}C$**

To allow for a direct comparison between different periods, in which the $\delta^{13}C$ of air may have been different, $\Delta^{13}C$ was calculated (following Farquhar et al. (1982)):

$$(\delta^{13}C_{air} - \delta^{13}C_{sample}) / (1 - (\delta^{13}C_{sample}/1000))$$

The value for $\delta^{13}C_{air}$ was derived from Ferrio et al. (2005), who based this on $\delta^{13}C$ values in laminated, and therefore well-dated, ice cores. For Sharara, we used the dates of 9250-9200 cal BCE, based on unpublished radiocarbon results, for the el-Hemmeh PPNA samples we used 9400-8700 cal BCE, and for LPPNB el-Hemmeh 7500-7000 BCE.

**Software and statistical analysis**

The results were processed using MS Excel and analysed as well as visualized using R Statistical Software (v. 4.1.2 and v.2024.04.2+764; R Core Team (2021)/(2024)) with the tidyverse, dplyr, ggplot, and viridis libraries. Statistical analyses were done using baseR and the packages rstatix and car. The results were checked for potential effects of the preservation status and presence of partial grains; there did not appear to be a consistent effect of either of these.

Shapiro-Wilks tests were used to confirm a normal distribution of the ($\Delta^{13}C$) data, in combination with visual inspection through density and QQ plots. According to the Shapiro-Wilks tests the $\Delta^{13}C$ data are normally distributed (although not the $\delta^{15}N$ data). Because the visual inspection appeared to show some skewedness nonetheless, Levene's test was used to check for homogeneity of variance, as it is less sensitive to deviations from normality. The variance was homogeneous for the analyses presented in this paper.

Differences between groups (sites, periods, domestication status) were assessed with unpaired 2-sample t-tests (Welch two sample t-tests) and one-way ANOVA tests. The Tukey post hoc tests was used to test which groups are different from each other. Linear correlation was assessed using a Spearman correlation test.

**Summary of results**

The full results are reported in Dataset S7 and have been published as a dataset in the Oxford University Research Archive [Oxford University Research Archive, DOI isotope dataset forthcoming]. Table S8 presents the average $\Delta^{13}C$ values for samples, Table S9 the average $\delta^{15}N$

values for samples. Table S10 and S11 present statistical results: Table S10 the results of the t-tests for $\Delta^{13}C$ values, directly comparing two independent groups, and Table S11 those of ANOVA and Tukey post hoc tests to dive deeper into the differences between domestication status groups.

For the nitrogen stable isotope results, it is worth noting that while some individual grains show very high values, the averages and the majority of the single-grain values are in line with the semi-arid region where the sites are, and typical for low-medium manuring input and values observed for wild plants and unmanured crops in the Southern Levant (Flohr, 2012; Hartman and Danin, 2010; Styring et al., 2016). The high variation and some very high values of individual grains is not unexpected; variation within a single site has been shown to be as large as 10‰ (Högberg, 1997; Lajtha and Marshall, 1994), although these studies were in colder areas; we have also observed such variation ourselves in modern fields in Jordan, (Flohr, 2012). For example, single plants may have grown in a dry, rocky spot, or in a marshy and/or particularly saline part of a field, with all of these conditions capable of producing high $\delta^{15}N$ in plants.

### Figures

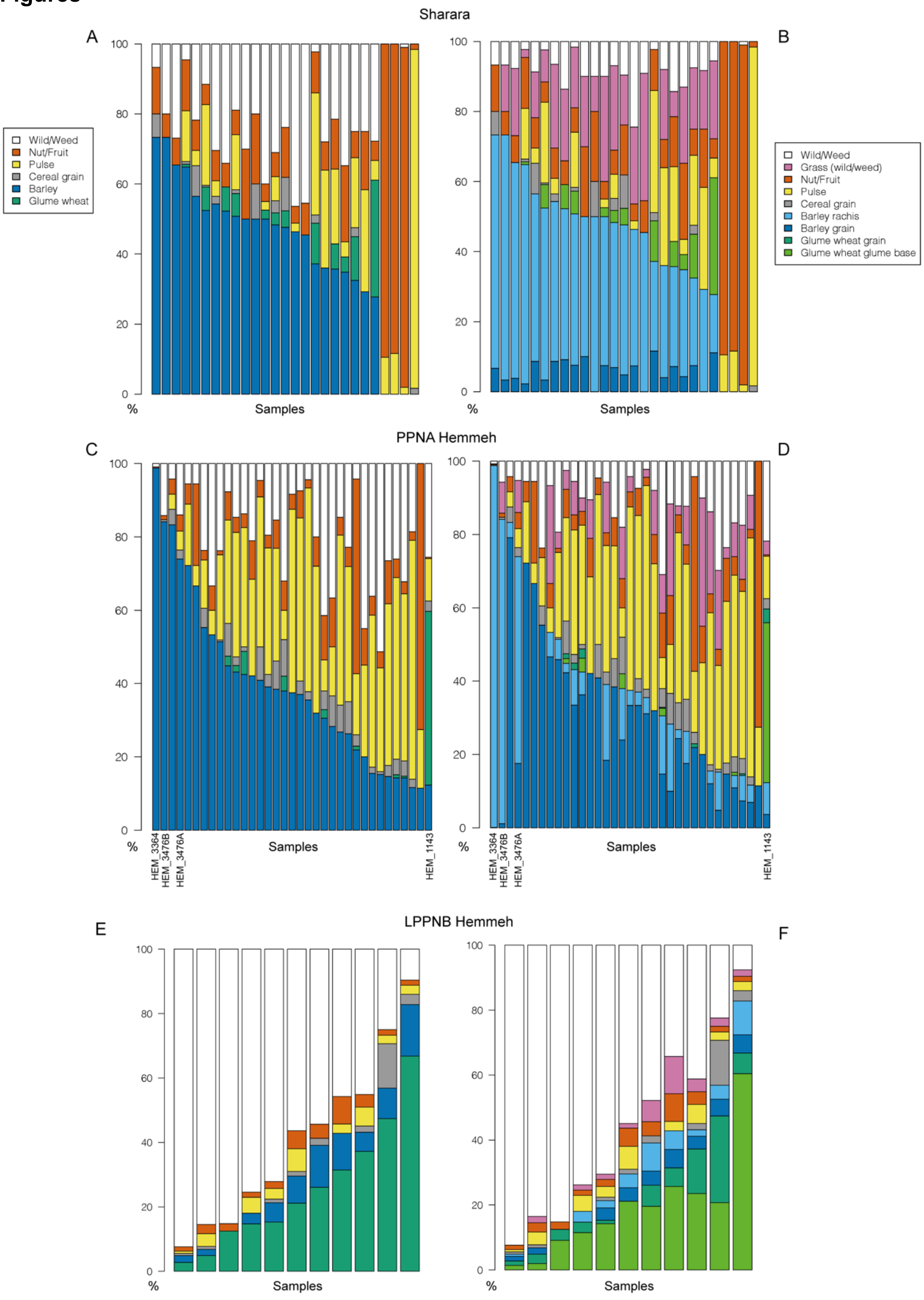
Sharara
A
B
Wild/Weed
Nut/Fruit
Pulse
Cereal grain
Barley
Glume wheat
Grass (wild/weed)
Barley rachis
Barley grain
Glume wheat grain
Glume wheat glume base
100
80
60
40
20
0
%
Samples
PPNA Hemmeh
C
D
HEM_3364
HEM_3476B
HEM_3476A
HEM_1143
LPPNB Hemmeh
E
F

**Fig. S1**. Relative abundance of major categories of plant remains across samples. (*A*) For Sharara, with graphs based on 'group 1' and (*B*) 'group 2' categories listed in Dataset S1. (*C*) At PPNA el-Hemmeh, with graphs based on 'group 1' and (*D*) 'group 2' categories listed in Dataset S2. (*E*) At LPPNB el-Hemmeh, with graphs based on 'group 1' and (*F*) 'group 2' categories listed in Dataset S2.

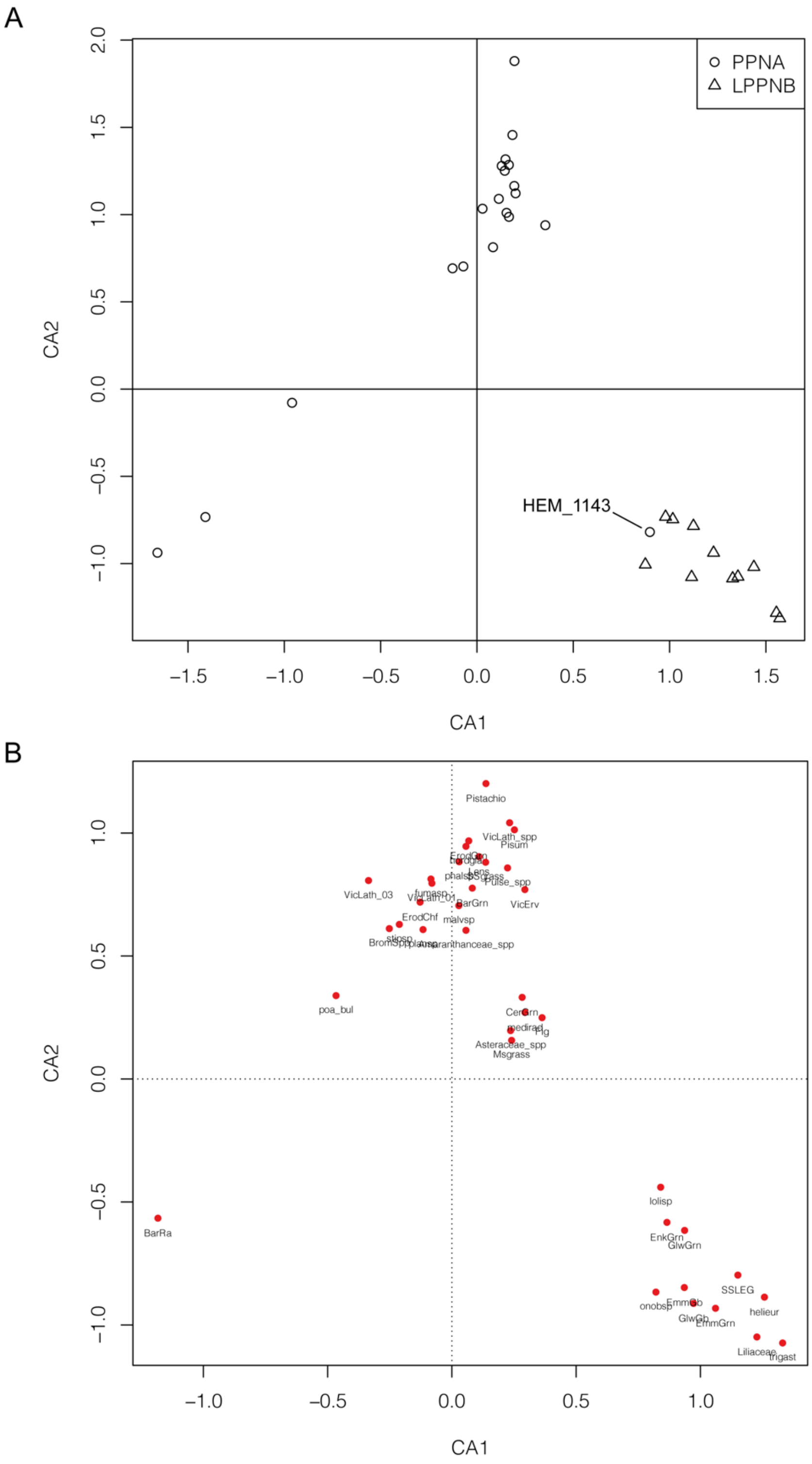

**Fig. S2**. Correspondence analysis plots of (*A*) 31 el-Hemmeh samples on the basis of 38 taxa, coded according to period (PPNA, LPPNB), and (*B*) 38 taxa based on their occurrence in 31 samples. Codes used for taxa in the analysis ('corro code') are reported in Dataset S2. A list of samples by period is provided in Dataset S3.

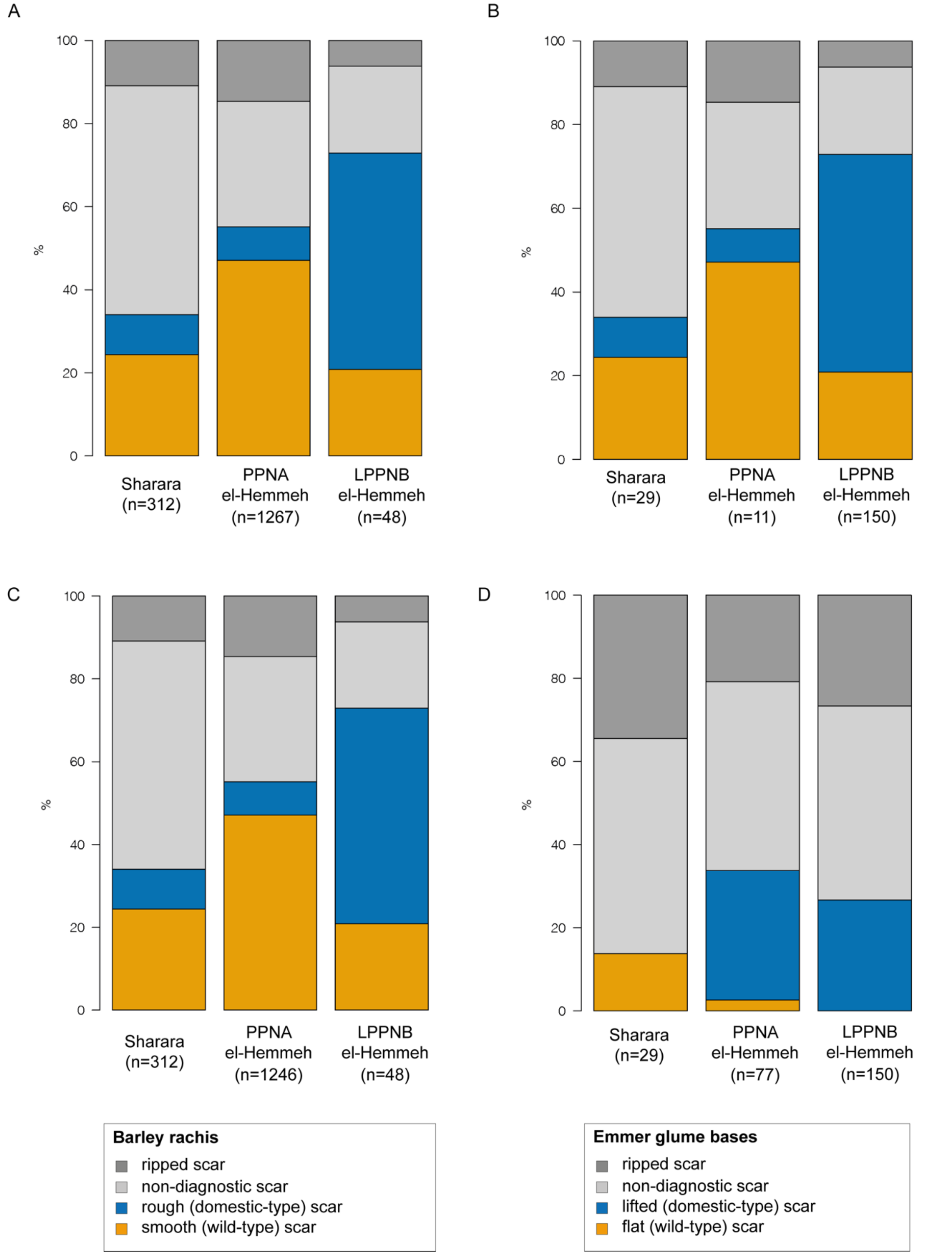

**Fig. S3**. Bar charts illustrating the relative percentage of different rachis morphotypes at Sharara, and both PPNA and LPPNB el-Hemmeh. (*A*) Barley. (*B*) Emmer. (*C*) Barley with sample HEM_1143 removed. (*D*) Emmer with sample HEM_1143 removed.

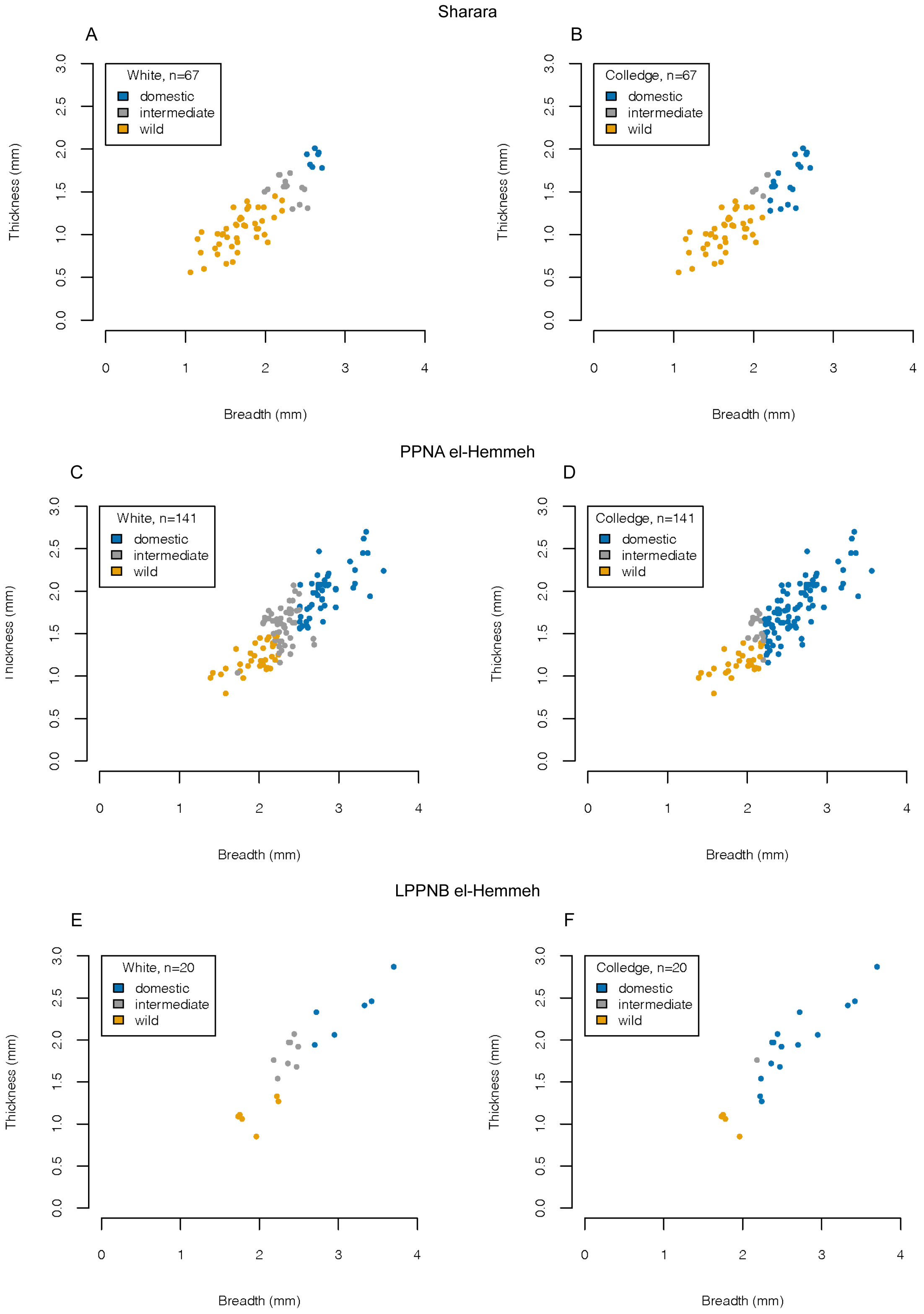

**Fig. S4**. Scatterplots showing breadth and thickness measurements of individual barley grains/grain fragments, coded according to whether these are categorised as wild, intermediate or domestic using the parameters published by White (2013, p. 96) and Colledge (2001, p. 64). (*A, B*) Sharara. (*C, D*) PPNA el-Hemmeh. (*E, F*) LPPNB el-Hemmeh.

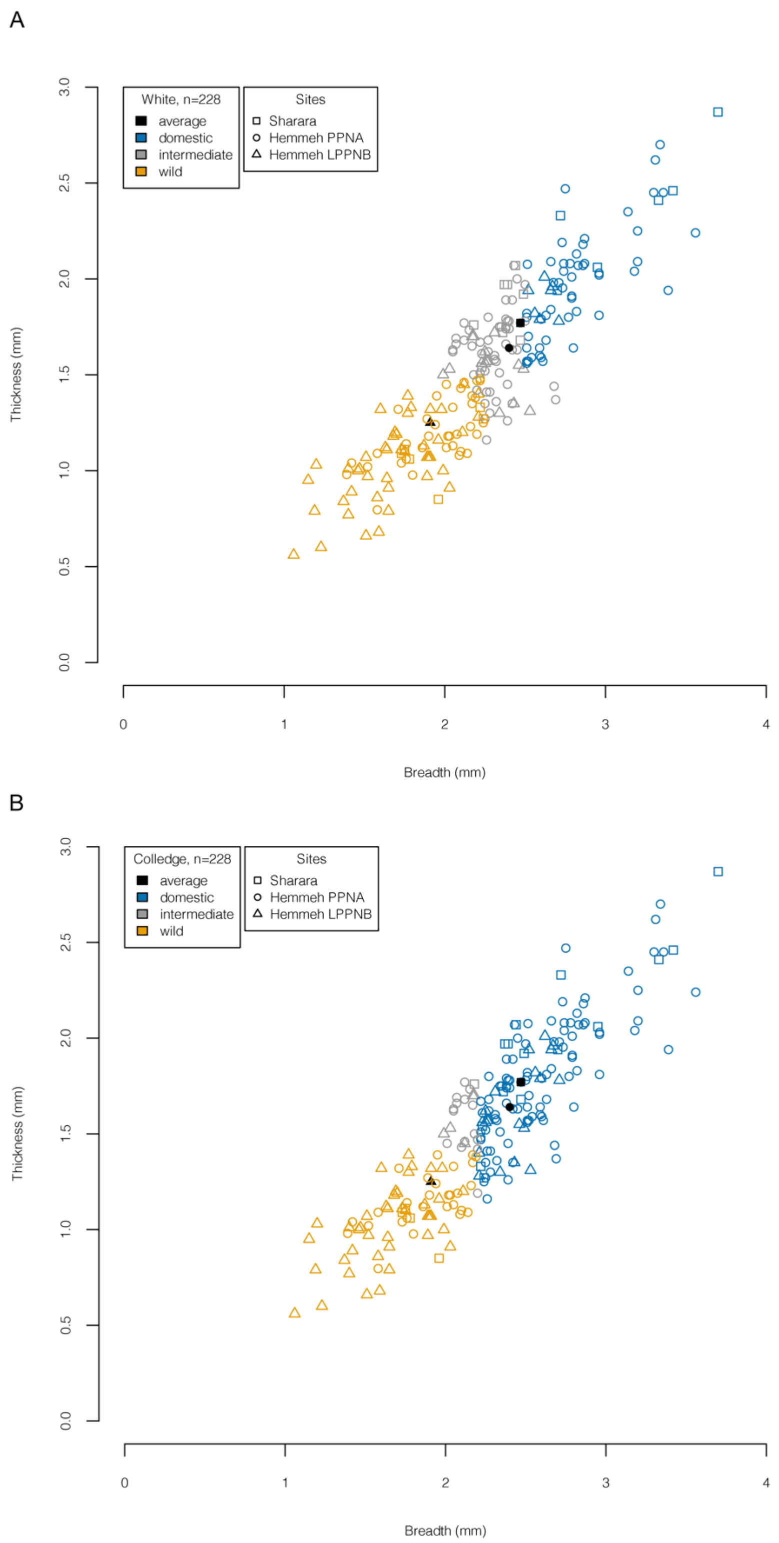

**Fig. S5**. Scatterplots showing breadth and thickness measurements of individual barley grains/grain fragments, coded by site and according to whether these are categorised as wild, intermediate or domestic. (*A*) Following White (2013, p. 96). (*B*) Following Colledge (2001, p. 64). Solid black symbols indicate average grain size calculated for each site.

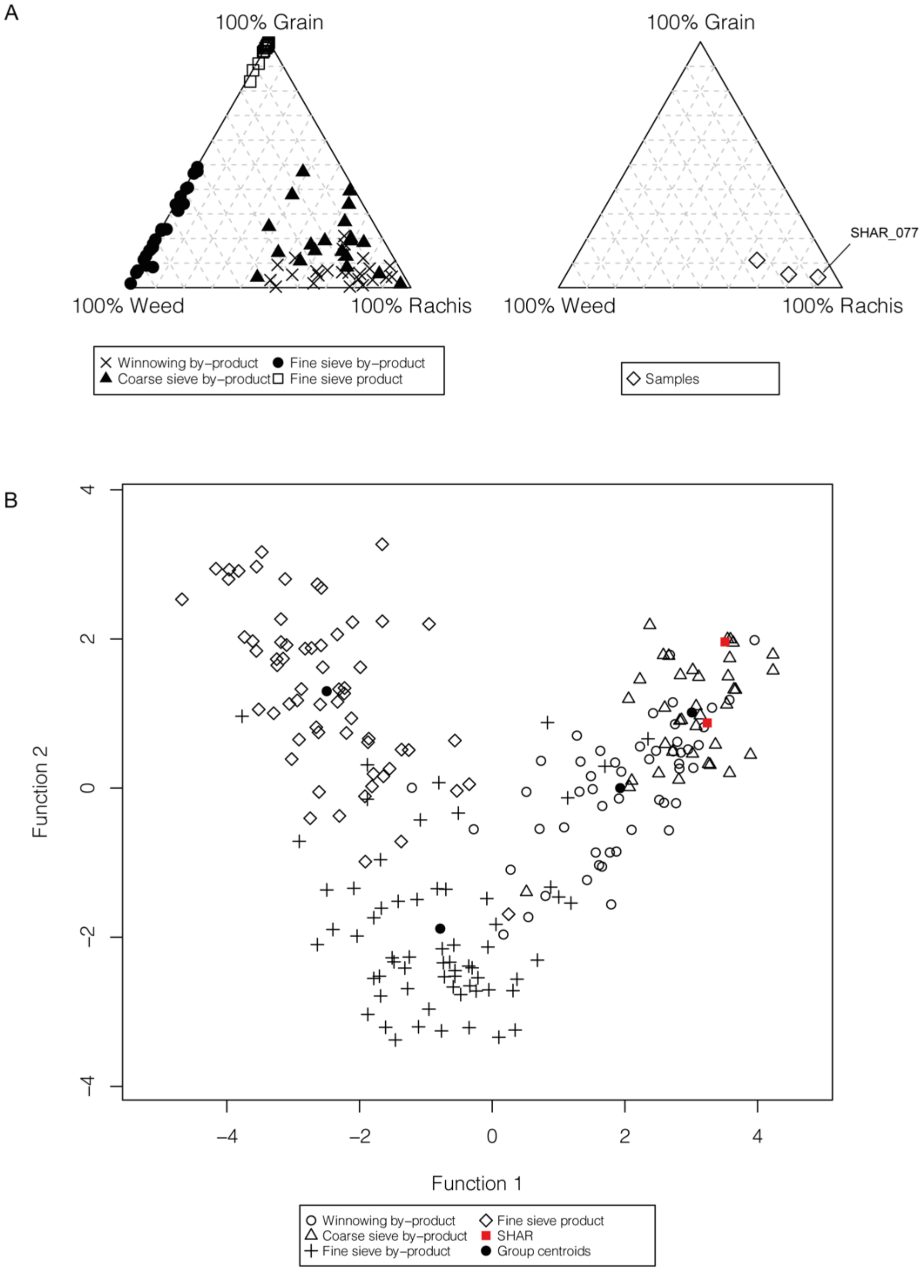

**Crop-processing classification**

| | | |
|---|---|---|
| SHAR_003 | Coarse-sieve by-product | Probability: 0.998 |
| SHAR_069 | Coarse-sieve by-product | Probability: 0.976 |

**Fig. S6**. Crop processing analysis on Sharara samples with at least 30 cereal items. (*A*) Triplot of three Sharara samples based on the relative frequency (%) of grain, rachis and weed items in samples, in comparison to ethnographic Amorgos samples. (*B*) Scatter plot of two archaeological samples from Sharara containing 10 or more weed items with definable physical properties in comparison to ethnographic samples from Amorgos on the first two discriminant functions (classification and probability of classification reported below).

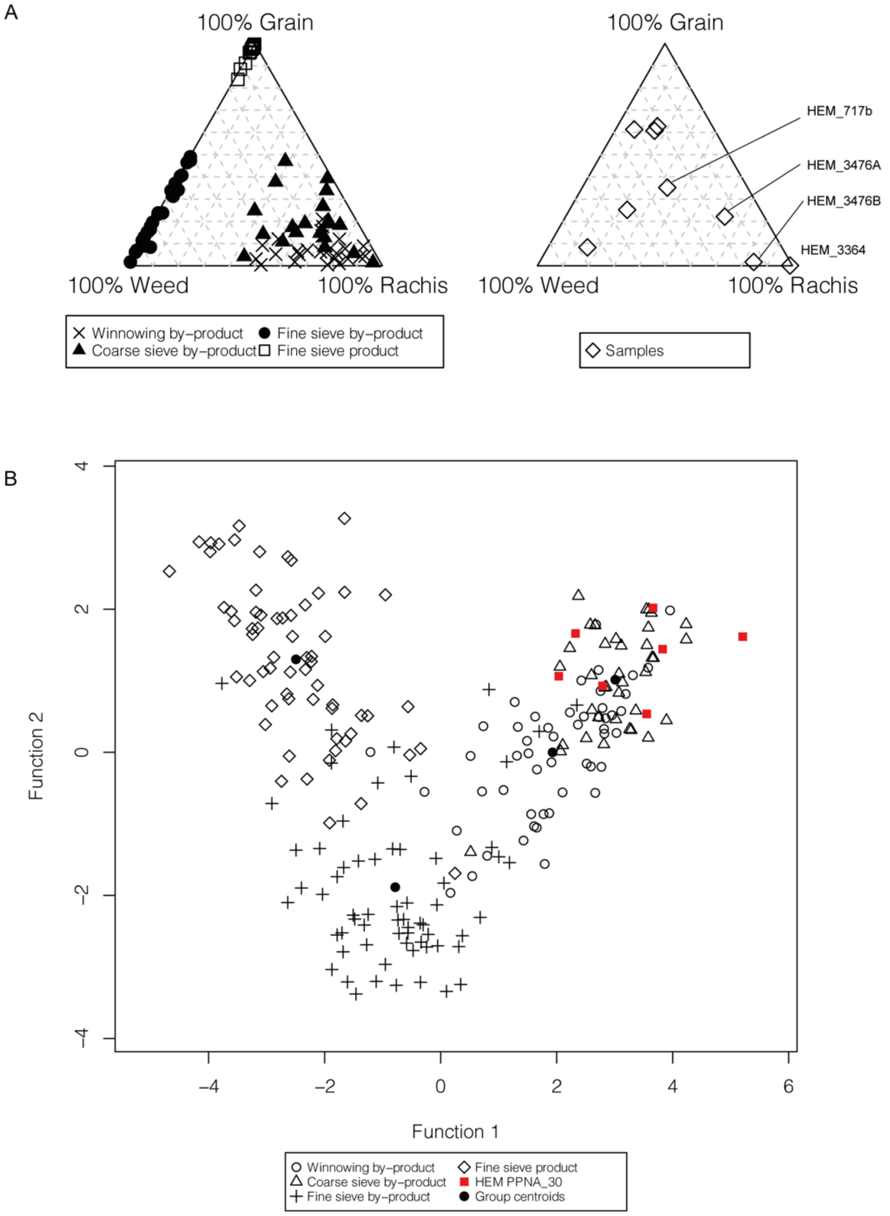

**Crop-processing classification**

| | | |
|---|---|---|
| HEM_3476A | Winnowing by-product | Probability: 0.999 |
| HEM_3476B | Coarse-sieve by-product | Probability: 0.863 |
| HEM_3237 | Coarse-sieve by-product | Probability: 0.977 |
| HEM_720 | Coarse-sieve by-product | Probability: 1.000 |
| HEM_815 | Coarse-sieve by-product | Probability: 0.998 |
| HEM_886 | Coarse-sieve by-product | Probability: 0.983 |
| HEM_721A | Coarse-sieve by-product | Probability: 1.000 |

**Fig. S7**. Crop processing analysis on PPNA el-Hemmeh samples with at least 30 cereal items. (*A*) Triplot of nine PPNA el-Hemmeh samples based on the relative frequency (%) of grain, rachis and weed items in samples, in comparison to ethnographic Amorgos samples. (*B*) Scatter plot of seven archaeological samples from PPNA el-Hemmeh containing 10 or more weed items with definable physical properties in comparison to ethnographic samples from Amorgos on the first two discriminant functions (classification and probability of classification reported below).

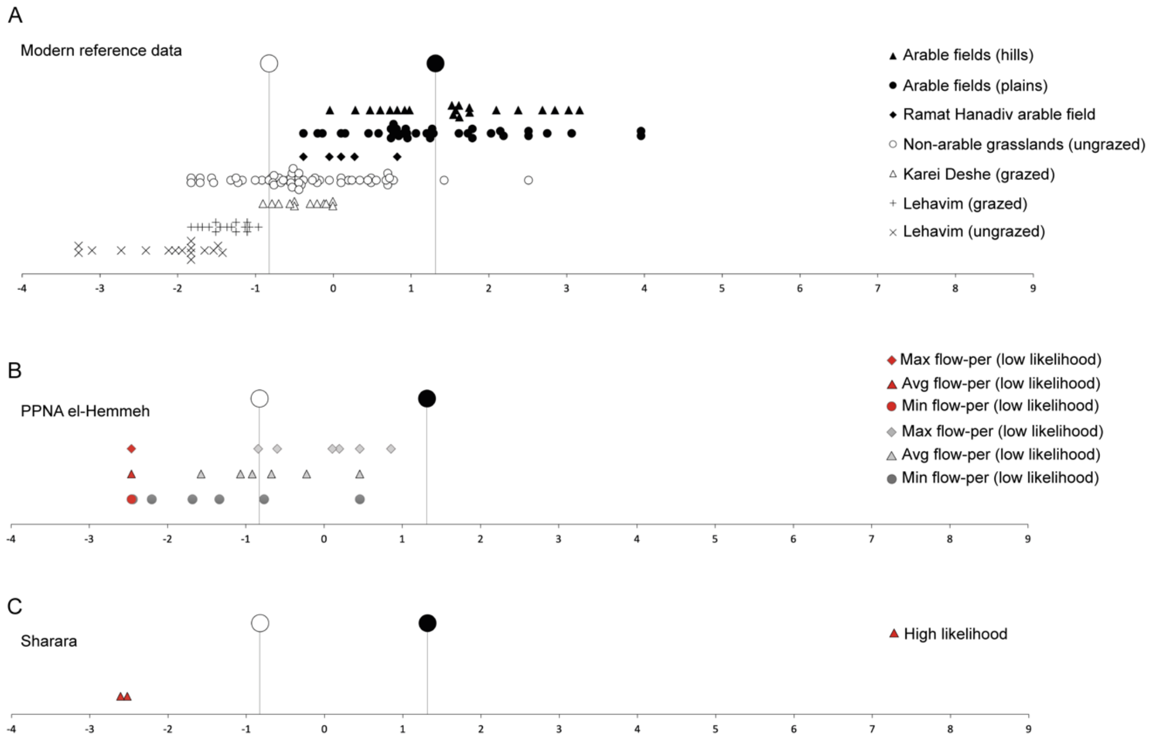

**Fig. S8.** Discriminant function extracted from the DA using only flowering duration as a discriminating variable. (*A*) Modern reference data from arable (closed symbols) and non-arable (open symbols) habitats; 86.7% of all modern samples were correctly reclassified (taken from (Weide et al., 2022)); the larger closed and open symbols that appear in all panels indicate group centroids. (*B*) Samples from PPNA el-Hemmeh according to whether average, minimum or maximum flowering duration was used for aggregate species with large flowering ranges, and whether samples had a high or low likelihood of representing crops and their associated weeds based on crop processing analysis. (*C*) Samples from Sharara, coded according to whether samples had a high or low likelihood of representing crops and their associated weeds based on crop processing analysis.

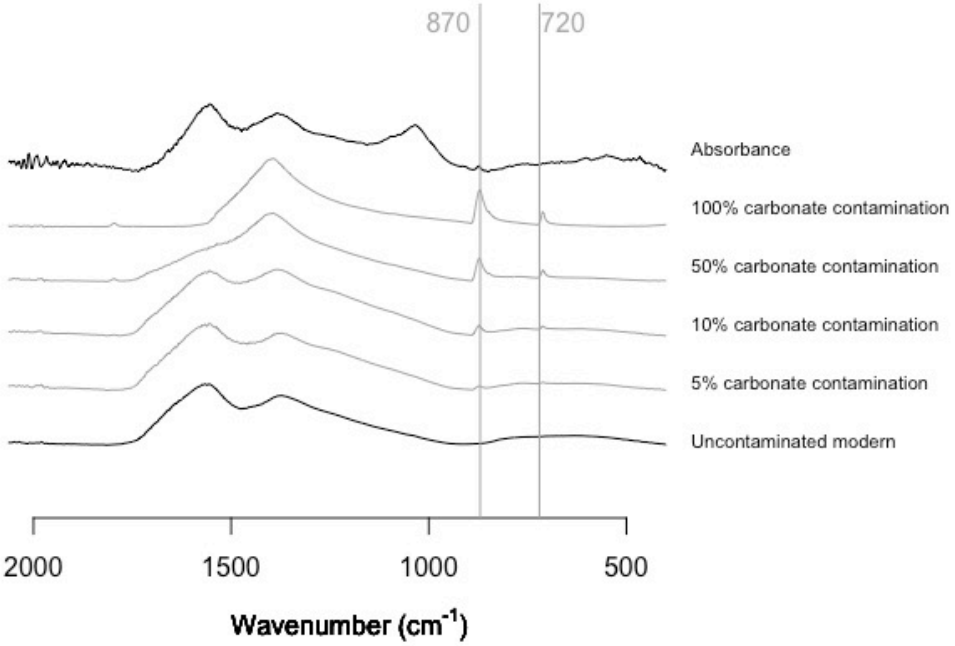

**Fig. S9.** Plot to compare the spectra of archaeological sample SHAR050A to those of modern grains with deliberate carbonate contamination (Vaiglova et al., 2014), reusing an R script written by Elizabeth Stroud. A slight carbonate peak is visible at 870 $cm^{-1}$ in the archaeological sample. Also note the humic peak to the left of the carbonate peak, likely due to sample-internal humics. Such plots were made for each of the FTIR-analysed samples with each of the potential contaminate. Please note the plots were made as a visual aid to the analysis and the aim was not to make them aesthetically pleasing.

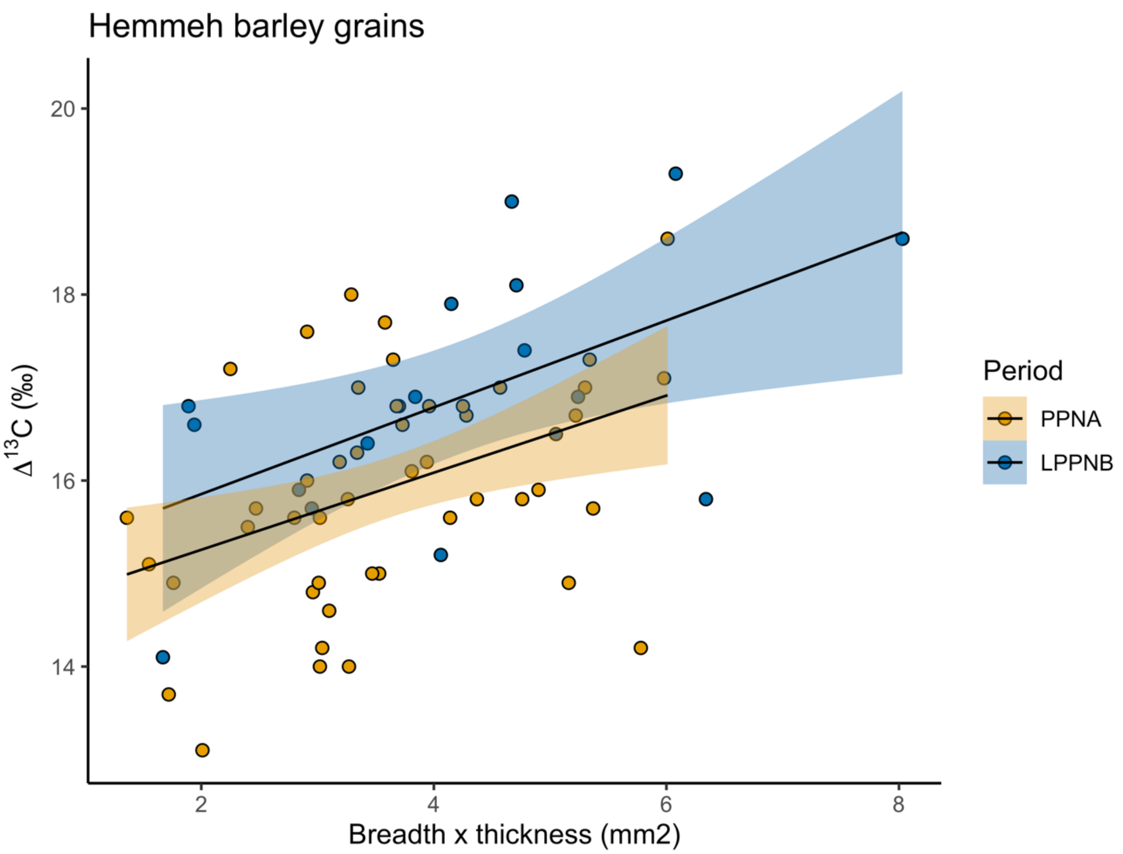

**Fig. S10.** Significant but weak correlation between size (breadth times thickness in $mm^2$) and $\Delta^{13}C$ for the el-Hemmeh barley grains. For this figure one outlier of a very large grain of over 10 $mm^2$ was removed from the LPPNB data. Yellow/orange/light = PPNA, blue/dark = LPPNB. Plotted using R Studio, ggplot library with the ‘geom_smooth’ function with the lm (linear model) method.

### Tables

**Table S1.** Summary of Sharara and PPNA el-Hemmeh samples included in weed ecological analysis that contained at least 100 cereal items, or at least 30 cereal items, and whether the samples are considered to have a high or low likelihood of representing crops and their associated weeds on the basis of crop processing analysis, as described in text and Tables S2 and S3.

| | **Weed ecology threshold** | | **Likelihood that sample represents crops and their associated weeds based on crop processing analysis** |
|---|---|---|---|
| **Sample** | **100 cereal items** | **30 cereal items** | |
| SHAR_003 | Included | Included | High |
| SHAR_069 | Included | Included | High |
| SHAR_077 | n/a | n/a | Low |
| HEM_3237 | Included | Included | Low |
| HEM_3476A | Included | Included | High |
| HEM_3476B | Included | Included | Low |
| HEM_720 | Included | Included | Low |
| HEM_815 | n/a | Included | Low |
| HEM_886 | n/a | Included | Low |
| HEM_721A | n/a | Included | Low |
| HEM_3364 | n/a | n/a | Low |
| HEM_717b | n/a | n/a | Low |

**Table S2.** Sharara samples included in crop processing analyses and the results of grain, rachis and weed proportions analysis (triplot) and weed-based discriminant analysis. If both approaches yielded consistent results, samples were considered to have a high likelihood or representing crops and their associated weeds, if not they were considered to have a low likelihood of representing crops and their associated weeds. *Sample which only met threshold for inclusion when lowered to 30 cereal items.

| Sample | In triplot | In DA | Likelihood |
|---|---|---|---|
| SHAR_003 | Winnowing by-product, Coarse sieve by-product | Coarse sieve by-product | High |
| SHAR_069 | Winnowing by-product, Coarse sieve by-product | Coarse sieve by-product | High |
| SHAR_077* | Winnowing by-product, Coarse sieve by-product | Not enough weed items | Low |

**Table S3.** PPNA el-Hemmeh samples included in crop processing analyses and the results of grain, rachis and weed proportions analysis (triplot) and weed-based discriminant analysis. If both approaches yielded consistent results, samples were considered to have a high likelihood or representing crops and their associated weeds, if not they were considered to have a low likelihood of representing crops and their associated weeds. * Sample which only met threshold for inclusion when lowered to 30 cereal items.

| Sample | In triplot | In DA | Likelihood |
|---|---|---|---|
| HEM_3237 | Ungrouped | Coarse sieve by-product | Low |
| HEM_3364 | Winnowing by-product, Coarse sieve by-product | Not enough weed items | Low |
| HEM_3476A | Winnowing by-product, Coarse sieve by-product | Winnowing by-product | High |
| HEM_3476B | Winnowing by-product, Coarse sieve by-product | Coarse sieve by-product (low probability) | Low |
| HEM_720 | Ungrouped | Coarse sieve by-product | Low |
| HEM_815* | Ungrouped | Coarse sieve by-product | Low |
| HEM_886* | Ungrouped | Coarse sieve by-product | Low |
| HEM_717b* | Coarse sieve by-product (borderline) | Not enough weed items | Low |
| HEM_721A* | Ungrouped | Coarse sieve by-product | Low |

**Table S4.** Identifications of wild/weed taxa at Sharara, their weed ecological code, flowering duration (flow_per), and for aggregate species groups a list of species included in each group. For taxa with aggregate species groups that had wider flowering duration ranges, minimum and maximum flowering durations for the aggregate species group are reported.

| Published identification | Weedeco code | flow_per | Aggregated species | Comments |
|---|---|---|---|---|
| cf. *Eremopyrum* sp. | eremsp | 2.5 | *Eremopyrum bonaepartis*<br>*Eremopyrum distans* | Two species of *Eremopyrum* are listed in the Flora Palaestina. These have comparable flowering durations, and this taxon has been included in the analyses as an aggregate group. |
| *Aegilops* cf. *speltoides* | aegispe | 3 | *Aegilops speltoides* | |
| *Aegilops* sp. | aegisp | 2.9 | *Aegilops bicornis*<br>*Aegilops sharonensis*<br>*Aegilops longissimi*<br>*Aegilops searsii*<br>*Aegilops peregrina*<br>*Aegilops kotschyi*<br>*Aegilops biuncialis*<br>*Aegilops geniculata*<br>*Aegilops crassa*<br>*Aegilops triuncialis* | Eleven species of *Aegilops* are listed in the Flora Palaestina. These have comparable flowering durations, and this taxon has been included in the analyses as an aggregate group after excluding *A. speltoides* on morphological grounds (see Whitlam et al. 2023; S2). |
| *Hordeum glaucum*<br>cf. *Hordeum glaucum* | hordgla | 3 | *Hordeum glaucum* | |
| *Taeniatherum critinum* | taencri | 3 | *Taeniatherum critinum* | |
| *Bromus* sp. | bromsp | 2.8 | *Bromus syriacus*<br>*Bromus tomentellus*<br>*Bromus brachystachys*<br>*Bromus Japonicus*<br>*Bromus danthoniae*<br>*Bromus lanceolatus*<br>*Bromus alopecuros*<br>*Bromus scoparius*<br>*Bromus tectorum*<br>*Bromus fasiculatus*<br>*Bromus rubens*<br>*Bromus madritensis*<br>*Bormus sterilis*<br>*Bromus diandrus*<br>*Bromus rigidus* | Sixteen species of *Bromus* are listed in the Flora Palaestina. One species (*B. catharticus*) is reported as a naturalised species first recorded in 1939. This was excluded. The remaining 15 species all have comparable flowering durations, and this taxon has been included in the analyses as an aggregate group. |
| *Avena* sp. | *avensp* | 2.2 | *Avena barbata*<br>*Avena wiestii*<br>*Avena clauda*<br>*Avena eriantha*<br>*Avena longiglumis* | Six species of *Avena* are listed in the Flora Palaestina. These have comparable flowering durations, and this taxon has been included in the analyses as an aggregate group after excluding *A. sterilis* on morphological grounds (Whitlam et al. 2023, S2). |
| *Poa bulbosa*<br>cf. *Poa bulbosa* | | 2 | *Poa bulbosa* | |
| *Poa* sp. | N/A | N/A | N/A | Seven species of *Poa* are listed in the Flora Palaestina. These species all have flowering durations that vary between one and seven months. As it was not possible to narrow the identification down to a single species, or group of species with comparable flowering durations, and as there were only two specimens of this type found at the site, this taxon was excluded from the analyses. |
| *Piptatherum holciforme/blancheanum* | *piptsp* | 3 | *Piptatherum holciforme/blancheanum* | |
| *Stipa capensis* | *stipcap* | 3 | *Stipa capensis* | |
| *Stipa* sp. | *stipsp* | 3.2 | *Stipa parviflora*<br>*Stipa capensis*<br>*Stipa barbata* | Six species of *Stipa* are listed in the Flora Paleastina. These have comparable flowering durations, and |

| | | | | |
|---|---|---|---|---|
| | | | *Stipa hohenackeriana*<br>*Stipa lagascae*<br>*Stipa bromoides* | this taxon has been included in the analyses as an aggregate group. |
| Medium-seeded Poaceae indeterminate | N/A | N/A | N/A | Excluded as not identified past family level. |
| *Capparis* sp. | N/A | N/A | N/A | Excluded as woody shrub. |
| *Brassica/Sinapis* sp. | brassp | 4 | *Brassica tournefortii*<br>*Brassica nigra*<br>*Sinapis alba*<br>*Sinapis arvensis* | Two species of *Brassica* and two species of *Sinapis* are listed in the Flora Palaestina. These have comparable flowering durations, and the taxon has been included in the analyses as an aggregate group |
| Brassicaceae sp. 'flat type' | N/A | N/A | N/A | Excluded as not identified past family level. |
| cf. Brassicaceae sp. | N/A | N/A | N/A | Excluded as not identified past family level. |
| *Erodium* sp. | erodsp | 2.7 | *Erodium hirtum*<br>*Erodium bryonifolium*<br>*Erodium deserti*<br>*Erodium subtrilobum*<br>*Erodium subintegrifolium*<br>*Erodium alnifolium* | Eighteen species of *Erodium* are listed in the Flora Palaestina. The specimens from Sharara correspond well with those from el-Hemmeh and the same species have been excluded as identifications. This leaves 6 species that have comparable flowering durations, and the taxon has been included in the analyses as an aggregate group. |
| Small-seeded legumes | N/A | N/A | N/A | Excluded as not identified past family level. |
| *Malva* sp. | malvsp | 3.7<br><br>Min = 2<br>Max = 5 | *Malva aegyptia*<br>*Malva sylvestris*<br>*Malva nicaeensis*<br>*Malva parviflora*<br>*Malva oxyloba*<br>*Malva neglecta* | Six species of *Malva* are listed in the Flora Palaestina. These have flowering durations that vary between 2 and 5 months and this this taxon has therefore been included using average, min and max flowering durations. However, no samples including this taxon made it into the final analysis. |
| Malvaceae sp. | N/A | N/A | N/A | Excluded as not identified past family level. |
| *Ammi majus* | | 3 | *Ammi majus* | This species is listed as having a variable flowering duration of 3 to 7 months, from (Mar-) Jun - Aug (-Sep). It was included in the analyses taking the Jun-Aug flowering duration in line with Weide et al. 2023 |
| Solanaceae sp. | N/A | N/A | N/A | Excluded as not identified past family level. |
| *Plantago* sp.<br>cf. *Plantago* sp. | | 6.5 | *Plantago lancelota/lagopus* | This was further identified as *Plantago lancelota/lagopus* following criteria used to identify the same type at el-Hemmeh (Seed cat). |
| *Centaurea* sp. | N/A | N/A | N/A | Nineteen species of *Centaurea* are listed in the Flora Palaestina. These have flowering durations that vary between 2 and 5 months. As it was not possible to narrow the identification down to a single species, or group of species with comparable flowering durations, and as there was only one specimen of this taxon, this was excluded from the analyses. |
| Asteraceae sp. 1 | N/A | N/A | N/A | Excluded as not identified past family level. |
| Asteraceae sp. 2 | N/A | N/A | N/A | Excluded as not identified past family level. |

**Table S5.** Identifications of wild/weed taxa at el-Hemmeh, their weed ecological code, flowering duration (flow_per), and for aggregate species groups a list of species included in each group. For taxa with aggregate species groups that had wider flowering duration ranges, minimum and maximum flowering durations for the aggregate species group are reported.

| Published identification | Weedeco code | flow_per | Aggregated species | Comments |
|---|---|---|---|---|
| *Eremopyrum bonaepartis/distans*<br>cf. *Eremopyrum bonaepartis/distans* | eremsp | 2.5 | *Eremopyrum bonaepartis*<br>*Eremopyrum distans* | |
| *Aegilops* sp. 1<br>cf. *Aegilops* sp. | aegisp | 2.9 | *Aegilops bicornis*<br>*Aegilops sharonensis*<br>*Aegilops longissimi*<br>*Aegilops searsii*<br>*Aegilops peregrina*<br>*Aegilops kotschyi*<br>*Aegilops biuncialis*<br>*Aegilops geniculata*<br>*Aegilops crassa*<br>*Aegilops triuncialis* | Eleven species of *Aegilops* are listed in the Flora Palaestina. These have comparable flowering durations, and these taxa have been included in the analyses as aggregate groups after excluding *A. speltoides* on morphological grounds. |
| *Aegilops/Triticum* sp. | N/A | N/A | N/A | Excluded as genus identification uncertain. |
| *Hordeum glaucum*<br>cf. *Hordeum glaucum* | hordgla | 3 | *Hordeum glaucum* | |
| *Taeniatherum critinum* | taencri | 3 | *Taeniatherum critinum* | |
| *Bromus tectorum* | bromtec | 3 | *Bromus tectorum* | |
| *Bromus* sp. 1<br>*Bromus* sp. 2 | bromspp | 3.1 | *Bromus syriacus*<br>*Bromus tomentellus*<br>*Bromus brachystachys*<br>*Bromus japonicus*<br>*Bromus danthoniae*<br>*Bromus lanceolatus*<br>*Bromus alopecuros*<br>*Bromus scoparius* | Sixteen species of *Bromus* are listed in the Flora Palaestina. One species (*B. catharticus*) is reported as a naturalised species first recorded in 1939. This was excluded, along with species listed in the section *Genea* that were ruled out on morphological grounds. The remaining eight species have comparable flowering durations and these taxa have been included in the analyses as aggregate groups. |
| *Bromus* sp. indet. | bromsp | 2.8 | *Bromus syriacus*<br>*Bromus tomentellus*<br>*Bromus brachystachys*<br>*Bromus Japonicus*<br>*Bromus danthoniae*<br>*Bromus lanceolatus*<br>*Bromus alopecuros*<br>*Bromus scoparius*<br>*Bromus tectorum*<br>*Bromus fasiculatus*<br>*Bromus rubens*<br>*Bromus madritensis*<br>*Bormus sterilis*<br>*Bromus diandrus*<br>*Bromus rigidus* | Sixteen species of *Bromus* are listed in the Flora Palaestina. One species (*B. catharticus*) is reported as a naturalised species first recorded in 1939. This was excluded. The remaining 15 species all have comparable flowering durations, and this taxon has been included in the analyses as an aggregate group. |
| *Avena* sp.<br>cf. *Avena* sp. | avensp | 2.2 | *Avena barbata*<br>*Avena wiestii*<br>*Avena clauda*<br>*Avena eriantha*<br>*Avena longiglumis* | Six species of *Avena* are listed in the Flora Palaestina. These have comparable flowering durations, and this taxon has been included in the analyses as an aggregate group after excluding *A. sterilis* on morphological grounds. |
| *Phalaris* sp.<br>cf. *Phalaris* sp. | phalsp | 4.6 | *Phalaris brachystachys*<br>*Phalaris minor*<br>*Phalaris paradoxa*<br>*Phalaris tuberosa*<br>*Phalaris canariensis* | Five species of *Phalaris* are listed in the Flora Palaestina. These have comparable flowering durations, and this taxon has been included in the analyses as an aggregate group. |
| *Lolium perenne/rigidum*<br>cf. *Lolium perenne/rigidum* | lolisp | 3.2 | *Lolium perenne/rigidum* | |

| | | | | |
|---|---|---|---|---|
| *Poa bulbosa* | poa_bul | 2 | *Poa bulbosa* | |
| *Poa/Phleum* sp. | N/A | N/A | N/A | Excluded as genus identification uncertain. |
| *Piptatherum holciforme/blancheanum* | piptsp | 3 | *Piptatherum holciforme/blancheanum* | |
| *Stipa capensis* | stipcap | 3 | *Stipa capensis* | |
| *Stipa barbata/lagascae* | | 3 | *Stipa barbata/lagascae* | |
| *Stipa* sp. | | 3 | *Stipa parviflora*<br>*Stipa capensis*<br>*Stipa barbata*<br>*Stipa hohenackeriana*<br>*Stipa lagascae*<br>*Stipa bromoides* | Six species of *Stipa* are listed in the Flora Palaestina. These have comparable flowering durations, and this taxon has been included in the analyses as an aggregate group. |
| Small-seeded grass sp. 1 | N/A | N/A | N/A | Excluded as not identified past family level. |
| Medium-seeded Poaceae indeterminate | N/A | N/A | N/A | Excluded as not identified past family level. |
| Small-seeded Poaceae indeterminate | N/A | N/A | N/A | Excluded as not identified past family level. |
| *Silene* sp. | silesp | 3<br><br>Min=1<br>Max=6 | *Silene italica*<br>*Silene longipetala*<br>*Silene swertiifolia*<br>*Silene vulgaris*<br>*Silene succulenta*<br>*Silene reinwardtii*<br>*Silene linearis*<br>*Silene modesta*<br>*Silene aegyptiaca*<br>*Silene fuscata*<br>*Silene rubella*<br>*Silene sedoides*<br>*Silene behen*<br>*Silene muscipula*<br>*Silene crassipes*<br>*Silene trinervis*<br>*Silene villosa*<br>*Silene damascena*<br>*Silene palaestina*<br>*Silene arabica*<br>*Silene vivianii*<br>*Silene nocturna*<br>*Silene gallica*<br>*Silene oxyodonta*<br>*Silene tridentata*<br>*Silene colorata*<br>*Silene aperala*<br>*Silene coniflora*<br>*Silene conoidea*<br>*Silene macrodonta*<br>*Silene physalodes*<br>*Silene grisea*<br>*Silene hussonii*<br>*Silene papillosa*<br>*Silene telavivensis* | Thirty-five species of *Silene* are listed in the Flora Palaestina, which vary in their flowering duration from 1 to 6 months.<br><br>This taxon has therefore been included in the analyses as an aggregate group with the flowering duration averaged (3 months), as well as tested with the minimum (1 month) and maximum (6 month) flowering duration. |
| *Vaccaria pyramidata* | vaccpyr | 4 | *Vaccaria pyramidata* | |
| Caryophyllaceae sp. | N/A | N/A | N/A | Excluded as not identified past family level. |
| *Atriplex* cf. *hastata* | atrihas | 6 | *Atriplex hastata* | |
| *Atriplex* cf. *rosea* | atriros | 5 | *Atriplex rosea* | |
| *Atriplex* sp. seed core | N/A | N/A | N/A | Excluded as not identified to species. Only 2. Species large flowering range. |
| Amaranthaceae sp. indet (surrounded by tissue)<br>Amaranthaceae sp. indet<br>Amaranthaceae seed core | N/A | N/A | N/A | Excluded as not identified past family level. |

| | | | | |
|---|---|---|---|---|
| *Adonis* sp. 1<br>*Adonis* sp. 2 | adonsp | 2.8 | *Adonis aleppica*<br>*Adonis aestivalis*<br>*Adonis cupaniana*<br>*Adonis denata*<br>*Adonis annua* | Five species of *Adonis* are listed in the Flora Palaestina. These have comparable flowering durations, and this taxon has been included in the analyses as an aggregate group. |
| *Papaver* spp. | papasp | 3 | *Papaver subpiriforme*<br>*Papaver carmeli*<br>*Papaver syriacum*<br>*Papaver humile*<br>*Papaver polytrichum*<br>*Papaver hybridum* | Seven species of *Papaver* are listed in the Flora Palaestina. These have comparable flowering durations, and this taxon has been included in the analyses as an aggregate group after excluding *P. argemone* on morphological grounds. |
| *Fumaria densiflora/parviflora*<br>*Fumaria* cf. *densiflora/parviflora* | fumasp | 4.5 | *Fumaria densiflora/parviflora* | |
| *Capparis* sp. | N/A | N/A | N/A | Excluded as woody shrub. |
| *Brassica/Sinapis* sp.<br>cf. *Brassica/Sinapis* sp. | brassp | 4 | *Brassica tournefortii*<br>*Brassica nigra*<br>*Sinapis alba*<br>*Sinapis arvensis* | Two species of *Brassica* and two species of *Sinapis* are listed in the Flora Palaestina. These have comparable flowering durations, and the taxon has been included in the analyses as an aggregate group |
| *Erodium* sp.<br>cf. *Erodium* sp. | erodsp | 2.7 | *Erodium hirtum*<br>*Erodium bryonifolium*<br>*Erodium deserti*<br>*Erodium subtrilobum*<br>*Erodium subintegrifolium*<br>*Erodium alnifolium* | Eighteen species of *Eremopyrum* are listed in the Flora Palaestina of which six are a possible match for the archaeological specimens based on size and morphological comparisons (Seed Catalogue). These have comparable flowering durations, and the taxon has been included in the analyses as an aggregate group. |
| *Scorpiurus muricatus* | scormur | 2 | *Scorpiurus muricatus* | |
| *Onobrychis* sp. 1<br>*Onobrychis* sp. 2 | onobsp | 2.3 | *Onobrychis supina*<br>*Onobrychis cadmea*<br>*Onobrychis kotschyana*<br>*Onobrychis ptolemaica*<br>*Onobrychis wettsteinii*<br>*Onobrychis caput-galli*<br>*Onobrychis crista-galli*<br>*Onobrychis squarrosa* | Eight species of *Onobrychis* are listed in the Flora Palaestina. These have comparable flowering durations, and these taxa have been included in the analyses as aggregate groups. |
| *Trigonella astroites* | trigsp | 2 | *Trigonella astroites* | |
| *Medicago radiata* | medirad | 3 | *Medicago radiata* | |
| Small-seeded legumes | N/A | N/A | N/A | Excluded as not identified past family level. |
| *Malva* sp. | malvsp | 3.7<br><br>Min=2<br>Max=5 | *Malva aegyptia*<br>*Malva sylvestris*<br>*Malva nicaeensis*<br>*Malva parviflora*<br>*Malva oxyloba*<br>*Malva neglecta* | Six species of *Malva* are listed in the Flora Palaestina. These have flowering durations that vary between 2 and 5 months and this this taxon has therefore been included in analysis 1 using average, min and max flowering durations. |
| Apiaceae sp. | N/A | N/A | N/A | Excluded as not identified past family level. |
| *Androsace maxima* | andrmax | 2 | *Androsace maxima* | |
| *Heliotropium* cf. *europaeum,* cf. *Heliotropium* cf. *europaeum* | helieur | 6 | *Heliotropium europaeum* | |
| cf. Lamiaceae sp. | N/A | N/A | N/A | Excluded as not identified past family level. |
| *Veronica* cf. *persica/polita* | verosp | 3.5 | *Veronica persica/polita* | |
| *Plantago* cf. *lancelota/lagopus* | plansp | 6.5 | *Plantago lancelota/lagopus* | |
| *Asperula arvensis* | aspearv | 2 | *Asperula arvensis* | |
| *Galium* cf. *aparine* | galiapa | 2 | *Galium aparine* | |

| | | | | |
|---|---|---|---|---|
| cf. *Galium* sp. | N/A | N/A | N/A | Twenty species of *Galium* are listed in the Flora Palaestina. These species have flowering durations that vary between 1 and 5 months. As it was not possible to narrow the identification down to a single species, or group of species with comparable flowering durations, and as there were only two specimens of this type found at the site, this taxon was excluded from the analyses. |
| *Centaurea* sp.<br>cf. *Centaurea* spp. | N/A | N/A | N/A | Nineteen species of *Centaurea* are listed in the Flora Palaestina. These have flowering durations that vary between 2 and 5 months. As it was not possible to narrow the identification down to a single species, or group of species with comparable flowering durations, and as there were only four specimens of this type identified at the site (with varying certainty), this taxon was excluded from the analyses. |
| *Anthemis* ‘type’ sp. | N/A | N/A | N/A | Excluded as genus identification uncertain. |
| *Leontondon* ‘type’ sp. | N/A | N/A | N/A | Excluded as genus identification uncertain. |
| Asteraceae sp. 1 | N/A | N/A | N/A | Excluded as not identified past family level. |
| Asteraceae sp. 2 | N/A | N/A | N/A | Excluded as not identified past family level. |
| Asteraceae sp. 3 | N/A | N/A | N/A | Excluded as not identified past family level. |
| Asteraceae/Dipsicaceae | N/A | N/A | N/A | Excluded as not identified past family level. |
| Liliaceae sp.<br>cf. Liliaceae sp. | N/A | N/A | N/A | Excluded as not identified past family level. |
| Cyperaceae sp. | N/A | N/A | N/A | Excluded as not identified past family level. |

**Table S6.** Isotope analysis sample sizes (individual grains) per site, period, and domestication status (following the White 2013 criteria). The numbers reported here refer to the samples that yielded results and are part of the reported analyses.

| | Sharara | el-Hemmeh | |
|---|---|---|---|
| | PPNA | PPNA | LPPNB |
| Barley | 32 | 48 | 18 |
| *-wild* | *13* | *16* | *5* |
| *-intermediate* | *14* | *18* | *8* |
| *-domestic* | *5* | *14* | *5* |
| Glume wheat | 0 | 10 | 35 |

**Table S7.** Standard reference materials used for calibration of $\delta^{13}C$ relative to VPDB (and $\delta^{15}N$ relative to AIR) and to monitor analytical uncertainty. The isotopic compositions of the in-house standards reported here represent long-term averages calibrated to VPDB and AIR with international standards like USGS40. The values used for calibrations and checks are in bold (e.g., LEU and P2 were only used for the nitrogen isotope calibrations).

| Standard | Material | Accepted /mean $\delta^{13}C$ (‰, VPDB) | Accepted / mean $\delta^{15}N$ (‰, AIR) |
|---|---|---|---|
| EMA-Spruce | Wood powder | **-25.44** | - 4.90 |
| EMA-Sorghum | Sorghum flour | **-13.78** | + 1.58 |
| ALA | Alanine | **-26.91** ±0.12 | **- 1.57** ±0.21 |
| SEAL | Seal bone collagen | **-12.60** ±0.11 | **+16.30** ±0.29 |
| COW | Cow bone collagen | **-24.28** | **+ 7.76** |
| LEU | Leucine | -23.23 | **+ 6.36** |
| EMA-P2 | Polymer | -28.19 ±0.14 | **- 1.57** ±0.19 |

**Table S8.** Average $\Delta^{13}C$ values in permille (‰) ± 1SD with sample size n in brackets. Domestication status is following the White 2013 criteria.

| | Sharara | el-Hemmeh | |
|---|---|---|---|
| | PPNA | PPNA | LPPNB |
| Barley | 17.0±1.4 (n=32) | 15.9±1.2 (n=48) | 16.8±1.4 (n=18) |
| -wild | 16.4±1.4 (n=13) | 15.5±1.3 (n=16) | 15.8±1.0 (n=5) |
| -intermediate | 17.5±1.3 (n=14) | 16.0±1.1 (n=18) | 17.2±1.2 (n=8) |
| -domestic | 17.1±1.5 (n=5) | 16.4±1.1 (n=14) | 17.3 ±1.5 (n=5) |
| Glume wheat | n/a | 17.4±1.4 (n=10) | 17.2±1.3 (n=35) |

**Table S9.** Average $\delta^{15}N$ values ± 1SD with sample size. Low sample sizes in red. Domestication status follows the White 2013 criteria.

| | Sharara | El-Hemmeh | |
|---|---|---|---|
| | PPNA | PPNA | LPPNB |
| Barley | 6.4±1.9 (n=7) | 6.8±2.4 (n=29) | 11.4±7.1 (n=3)* |
| -wild | 6.6±2.0 (n=6) | 7.2±2.3 (n=7) | n/a |
| -intermediate | 5.4 (n=1) | 6.2±2.8 (n=13) | 8.8 (n=1) |
| -domestic | n/a | 7.2±1.9 (n=9) | 12.7±9.5 (n=2)* |
| Glume wheat | n/a | 6.3±1.7 (n=4) | 6.1±2.2 (n=11) |

**including a high outlier*

**Table S10.** Results of two-sided t-tests for $\Delta^{13}C$ values, directly comparing two independent groups (Shapiro-Wilks normality tests conducted beforehand). Bold indicates p-values are significant at $p<0.05$. ANOVA analyses gave similar results (equal in terms of significant / non-significant).

| Group 1 | Group 2 | P-value | t | df |
|---|---|---|---|---|
| Sharara (=PPNA) barley | Hemmeh PPNA barley | **0.0008814** | 3.5023 | 59.57 |
| Sharara (=PPNA) barley | Hemmeh LPPNB barley | 0.714 | 0.36933 | 36.386 |
| Hemmeh PPNA barley | Hemmeh LPPNB barley | **0.01842** | -2.5046 | 27.706 |
| Hemmeh PPNA barley | Hemmeh PPNA wheat* | **0.007925** | -3.1811 | 11.973 |
| Hemmeh LPPNB barley | Hemmeh LPPNB wheat | 0.3159 | -1.0179 | 34.038 |

**note that only 10 glume wheat grain values were available for the el-Hemmeh PPNA.*

**Table S11a**. ANOVA results for $\Delta^{13}C$ and domestication status. Statistically significant in bold.

| Site | Domestication status criteria | Df (res) | SumSq | MS | F | p-value |
|---|---|---|---|---|---|---|
| El-Hemmeh | White 2013 | 2 (63) | 12.40 (98.02) | 6.20 (1.56) | 3.984 | **0.024** |
| El-Hemmeh | Colledge 2001 | 2 (63) | 14.98 (95.44) | 7.49 (1.52) | 4.945 | **0.010** |
| Sharara | White 2013 | 2 (29) | 8.85 (52.05) | 4.42 (1.80) | 2.464 | 0.103 |
| Sharara | Colledge 2001 | 2 (29) | 5.63 (55.27) | 2.81 (1.91) | 1.476 | 0.245 |

**Table S11b**. Tukey post hoc tests after ANOVA presented in Table S11a. Statistically significant in bold.

| | White 2013 | Colledge 2001 |
|---|---|---|
| *El-Hemmeh* | p-value | p-value |
| Wild – Intermediate | 0.106 | 0.069 |
| Wild – Domesticated | **0.023** | **0.009** |
| Intermediate – Domesticated | 0.679 | 0.964 |
| *Sharara* | | |
| Wild – Intermediate | 0.087 | 0.291 |
| Wild – Domesticated | 0.532 | 0.368 |
| Intermediate – Domesticated | 0.854 | 0.844 |

**Legends for Datasets**

**Dataset S1.** Species-by-sample data for 34 Sharara samples. Group 1 = major botanical categories, Group 2 = major botanical categories with cereals and wild/weed taxa further subdivided, Rachis ID = scar types as described in Supplementary Information and Main Text, corro code = codes used in correspondence analysis (not done for Sharara), weed eco code = codes used in weed ecological analysis, triplot = codes used to organise data to calculate percentages of grain, rachis and weed within samples, crop pro code = code used in weed-based discriminant analysis.

**Dataset S2.** Species-by-sample data for 46 el-Hemmeh samples. Group 1 = major botanical categories, Group 2 = major botanical categories with cereals and wild/weed taxa further subdivided, Rachis ID = scar types as described in Supplementary Information and Main Text, corro code = codes used in correspondence analysis, weed eco code = codes used in weed ecological analysis, triplot = codes used to organise data to calculate percentages of grain, rachis and weed within samples, crop pro code = code used in weed-based discriminant analysis.

**Dataset S3.** List of 46 el-Hemmeh samples and whether these are from PPNA or LPPNB levels.

**Dataset S4.** Breadth (mm) and thickness (mm) of 67 Sharara, 141 PPNA el-Hemmeh and 20 LPPNB el-Hemmeh barley grains/grain fragments, and their classification according to parameters published by White (2013, p. 96) and Colledge (2001, p. 64). Average grain size also included.

**Dataset S5.** List of species and summary of relevant information taken from the Flora Palaestina to inform weed ecological analysis. Flowering duration of species and calculation of flowering duration calculated as number of months.

**Dataset S6.** Average attribute scores for flowering duration (flow_dur), and discriminant scores (DS) and predicted group of archaeobotanical samples.

**Dataset S7.** Overview of analysed samples and results for the carbon stable isotope analyses.
Sample_name: Sample name consisting of the site abbreviation ('SHAR' for Sharara, ‘HEM’ for el-Hemmeh), the botanical sample number in three digits (e.g., '037'), and a letter to indicate different isotope samples taken from the botanical sample (e.g., 'SHAR069A', 'SHAR069B',
Site: Abbreviation for the site, SHAR for Sharara.
Period: The cultural period. PPNA = Pre-Pottery Neolithic A.
Locus: The archaeological context number, assigned by the excavators.
Botanical_sample: The archaeobotanical sample number, normally one sample per locus.
Grain: For ease of analysis, the grain indication also present in the sample_name is repeated here.
Species: Barley of GLW (=glume wheat) (technically not as precise as the species).
ID_White: Domestication status according to the metrics by White. Values: Domestic, Intermediate, Wild. See related paper of Whitlam et al. for an explanation.
ID_Colledge: Domestication status according to the metrics by Colledge. Values: Domestic, Intermediate, Wild.
Grain part: The part of the grain that was available for analysis.
Preservation: Preservation or charring state.
Weight: In mg.
B x T: Breadth times thickness.
d13C_av: Average carbon stable isotope ratio d13C in permille relative to VPDB.
d13C_sd: The standard deviation of the carbon stable isotope ratio if more than one measurement was taken.
d13C_charcorr: The carbon stable isotope ratio d13C after correction for charring (see methodological information).

d13Cair: The carbon stable isotope ratio d13C of the air, estimated for the period based on Ferrio et al. 2005.
D13C: Carbon stable isotope discrimination, calculated from the charring-corrected d13C and d13Cair (see methodological information).
